# Cl^-^-dependent amplification of excitatory synaptic potentials at distal dendrites revealed by voltage imaging

**DOI:** 10.1101/2023.05.29.542696

**Authors:** Masato Morita, Reo Higashi, Shin-ya Kawaguchi

## Abstract

Processing of synaptic signals in somatodendritic compartments determines the neuronal computation. Although amplification of excitatory signals by local voltage-dependent cation channels has been extensively studied, its spatio-temporal dynamics in elaborate dendritic branches remains obscure because of technical limitation. Using fluorescent voltage imaging throughout dendritic arborizations in hippocampal pyramidal neurons, here we demonstrate a unique Cl^-^-dependent remote computation mechanism equipped in distal branches. Local laser photolysis of caged-glutamate triggered excitatory postsynaptic potentials spreading along dendrites with gradual amplification toward the distal end whereas with attenuation toward the soma. *Tour-de-force* subcellular patch-clamp recordings from thin branches complemented with biophysical model simulation revealed that the asymmetric augmentation of excitation relies on the TTX-resistant Na^+^ channels and Cl^-^-conductances accompanied with deeper dendritic resting potential. Taken together, the present study unveils cooperative voltage-dependent actions of cation and anion conductances for dendritic supralinear computation which can locally decode the spatio-temporal context of synaptic inputs.

## Introduction

Neurons extend elaborate dendrites to receive synaptic inputs from other neurons. These synaptic inputs are spatially and temporally integrated in an analogue manner, and then the signals are transformed into action potentials (APs) as digital all-or-none signals at the axon initial segment. It has been shown that hippocampal and cortical pyramidal neurons exhibit several types of regenerative local spikes upon synaptic inputs in dendrites, including Na^+^, NMDA, and Ca^2+^ spikes (1, 2). Na^+^ and NMDA spikes are triggered by local multiple synaptic inputs in dendritic branches (3–6). On the other hand, Ca^2+^ spikes mediated by voltage-gated Ca^2+^ channels, show larger and broader waveforms, often leading to burst APs (1, 7, 8). In contrast to such a dendritic supra-linear computation mediated by active conductances, sub-threshold potential changes are thought to obey the passive membrane properties of dendrites, attenuating during the propagation as the cable theory predicts (9, 10). However, at present, it remains elusive how the subthreshold potential changes are processed in dendrite because of the difficulty to apply the patch-clamp technique to multiple points of a thin dendrite at the same time. As a corollary, mathematical model of electrical circuit assuming multiple compartments has provided basic view of signal integration in dendrites (11, 12), substantial part of which still remains to be consolidated by direct experimental analysis.

In this decade, genetically-encoded voltage indicators (GEVIs), which detect membrane potential changes as fluorescence changes, have drastically developed (13–15). This technique has enabled to fluorescently detect membrane potential changes at a tiny structure which is difficult to apply the patch-clamp technique, for example even a dendritic spine and a presynaptic bouton (16, 17). Taking advantage of the leading-edge voltage imaging technique, in this study, we studied how subthreshold synaptic potentials are processed in dendrites. Using improved version of ASAP1 (Accelerated Sensor of Action Potentials 1) (13), a fluorescent voltage imaging probe, expressed in a cultured hippocampal neuron in combination with very local glutamate or GABA stimulation on a dendrite by 405 nm spot-laser-mediated photoactivation, we found that even small size of glutamatergic excitatory postsynaptic potentials (EPSPs) spread in dendrites with opposite modulation depending on the direction of propagation. We also identified a novel Cl^-^-dependent mechanism underlying the unique asymmetric processing of the electrical signal in dendrites.

## Results

### Fluorescent imaging of membrane potential changes in a cultured hippocampal neuron

In this study, for better fluorescent detection of membrane potential changes in neurons, some amino acid replacements reported in an improved version ASAP3 (18) were introduced into the original ASAP1 (13) (here we note this version ASAP3β; for details, see methods). We evaluated fluorescence change of ASAP3β upon voltage changes by expressing in HEK293T cells (fig. S1A). ASAP3β exhibited fluorescence decrease (−40.0 ± 1.1 % ΔF/F, fig. S1, B and C) in response to 100 mV depolarization of the voltage-clamped HEK293T cell from a holding potential at −70 mV, which corresponded to almost two-fold sensitivity of the original ASAP1 (−18.5 ± 1.2 % ΔF/F, fig. S1C) similarly to ASAP3 (18).

We then examined how ASAP3β detects neuronal activity characterized by rapid membrane potential changes in a cultured hippocampal neuron by expressing it with electroporation (Fig. 1A). Both the fluorescence and the membrane potential were simultaneously recorded by imaging and patch-clamp methods. ASAP3β detected spontaneous EPSPs almost perfectly without any noticeable change in time course (Fig. 1B), showing ∼ 1.3 % fluorescence change for 2 mV potential change around −70 mV of membrane potential (Fig. 1C). On the other hand, more rapid spontaneous or evoked APs were detected as smaller fluorescence changes relative to the actual change of membrane potential (−8.4 ± 0.4 % ΔF/F upon 93.6 ± 2.7 mV, Fig. 1, B and C), likely because of the insufficient velocity of ASAP3β to follow a given rapid voltage change (Fig. 1, B and C). From these observations, it was confirmed that ASAP3β nicely detects synaptic potentials.

**Fig. 1.**
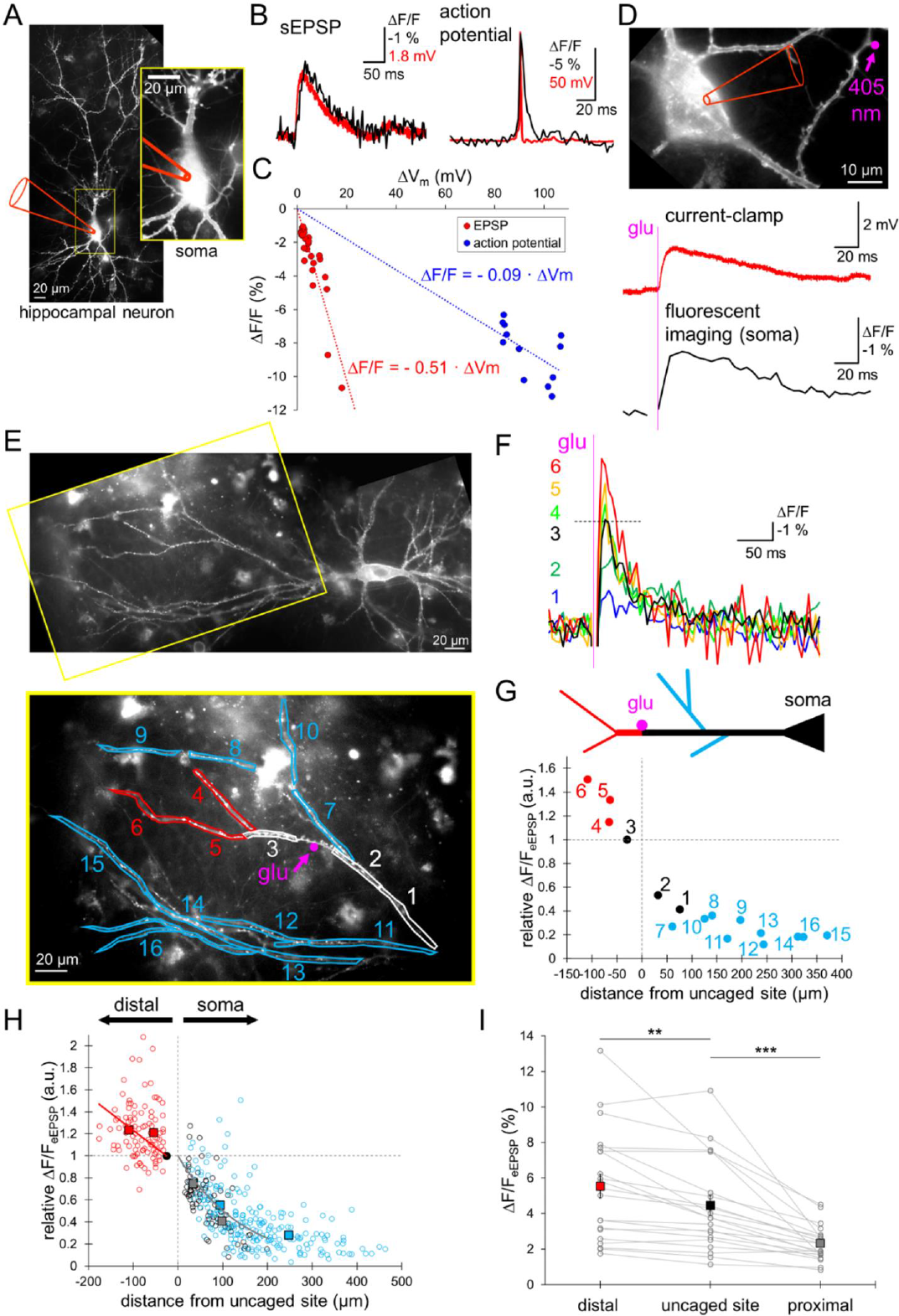
Direction-dependent EPSP modulation in dendrites by voltage imaging. **(A)** Fluorescent image of a cultured hippocampal neuron expressing ASAP3β under the whole-cell patch clamp recording. **(B)** Representative spontaneous EPSP (left) and action potential (right) in a cultured hippocampal neuron, recorded by ASAP imaging (black) and current-clamp recording (red). **(C)** Relative fluorescence changes (ΔF/F) for EPSPs (red, 29 events, 5 cells) and action potentials (blue, 12 events, 3 cells) plotted against the size of membrane potential changes (ΔV_m_). **(D)** Top, single-photon (405 nm) glutamate uncaging around a spine of an ASAP3β-expressing neuron under the whole-cell patch-clamp recording. Bottom, somatic voltage change upon glutamate uncaging recorded by current-clamp (red) or fluorescence imaging (black). **(E)** Top, image of an ASAP3β expressing neuron. Yellow rectangular area is expanded below, showing branches used for voltage imaging of local glutamate-evoked EPSPs. Magenta point indicates the location of 405 nm spot laser illumination. **(F)** ΔF/F_eEPSP_ traces at six ROIs (1–6) indicated in E. **(G)** Top, Schematic drawing of classification of dendrites (black, dendrite on the way to soma; red, distal branches; blue, bifurcated branches on the way to soma). Bottom, ΔF/F_eEPSP_ sizes for individual eEPSPs (normalized to that at ROI 3) in a cell (show in E) are plotted against the distance from the uncaged site. **(H)** Relative ΔF/F_eEPSP_ against the distance from the uncaged site (21 cells, red: 102, black: 85, blue: 226 regions based on the categorization shown in G). Square plots indicate averages (± SEM). **(I)** Absolute ΔF/F_eEPSP_ sizes at three categories of location: distal branch, uncaged site, and proximal branch (n = 21 cells, **: P < 0.01, ***: P < 0.001, paired t-test). Square plots indicate averages (± SEM). See also fig. S1 and S2.

### Direction-dependent EPSP modulation in dendrites

Taking advantage of real-time fluorescent detection of local membrane potential changes of interest throughout dendritic arborization, we studied how synaptic potentials propagate in dendrites. Glutamatergic EPSPs (EPSP_glu_) were evoked by local photo-activation of MNI-caged-glutamate (300 μM in the external bath) by 405 nm laser illumination (2 μm diameter) at a spine of ASAP3β-expressing hippocampal pyramidal neuron in culture (Fig. 1D). Simultaneous recordings of membrane potentials by current-clamp and fluorescent imaging from an ASAP3β-expressing neuron showed similar waveforms of EPSPs characterized by 1.4 ± 0.1 % (21 events) fluorescence change at the soma upon 2 mV potential change without temporal lag (Fig. 1D). Thus, evoked EPSP (eEPSP) measurement by ASAP3β enabled us to quantify the spatial pattern of EPSP spreading in a neuron.

Using this method, eEPSPs were monitored as the relative ASAP fluorescence changes (ΔF/F) throughout the dendrites of ASAP3β-expressing hippocampal neurons in the view field (Fig. 1, E and F). To focus on the propagation of EPSPs far from activation of typical voltage-gated active conductances, such as Na^+^ and Ca^2+^ channels, the laser intensity and duration was adjusted so that the EPSP size was around 2 - 5 mV (∼ 1 - 3 % of ΔF/F in ASAP fluorescence change). In line with the prediction of the cable theory (11), the fluorescence change for eEPSP (ΔF/F_eEPSP_) attenuated as propagation toward the soma (Fig. 1, E and G). The ΔF/F_eEPSP_ decreased by 25 ± 2 % at 33.6 ± 3.1 μm away from the uncaged site and by 59 ± 3 % at 97.7 ± 4.9 μm (Fig. 1H, black circles). Thus, the propagation of EPSPs toward the soma attenuates almost exponentially as characterized by a length constant of ∼ 125 μm. In contrast, surprisingly, EPSPs detected as ΔF/F_eEPSP_ rather amplified (in some cases to almost double, although variable), when propagating toward the distal end of a dendritic branch (Fig. 1, E to H and fig. S1D). The ΔF/F_eEPSP_ increased by 21 ± 4 % in average at 54.7 ± 1.9 μm away from the uncaged site and by 23 ± 3 % at 110.6 ± 4.5 μm (Fig. 1H, red). This is quite contrasting to the classical view of passive traveling of subthreshold membrane potential changes in a cable, in which membrane potential change attenuates during propagation in a dendritic tree regardless of direction. The ΔF/F_eEPSP_ amplification detected by ASAP imaging during propagation toward the distal end was not dependent on the absolute size of ΔF/F_eEPSP_ at the uncaged site (ranging from 1.1 % to 10.9 %) that reflects the magnitude of EPSPs (Fig. 1I and fig. S1D). Thus, our data indicated that excitatory synaptic inputs are oppositely modulated in a dendrite depending on the direction of propagation.

### GABAergic PSPs attenuate as spread in dendrites irrespective of direction

To further study the spatial pattern of spreading of membrane potential changes in a dendrite, we examined how GABAergic postsynaptic potentials (PSP_GABA_s) spread in dendrites by the ASAP3β imaging coupled with local activation of GABA_A_ receptors (GABA_A_Rs). PSP_GABA_s were caused by local uncaging of DPNI-caged-GABA (200-500 μM) with a spot 405 nm laser illumination (Fig. 2 and fig. S3), in a similar manner to the local glutamate stimulation. In many cases, local GABA stimulation on a dendrite of hippocampal neuron exhibited little fluorescence change of ASAP3β, reflecting the resting membrane potential (Vm_rest_) substantially close to the equilibrium potential for Cl^-^ (E_Cl_). Still, some cells exhibited clear increase of ASAP3β fluorescence upon GABA uncaging, corresponding to hyperpolarization (Fig. 2A and fig. S3A). In contrast, only very limited cases showed depolarization as observed with fluorescence decrease upon the local GABA, which is probably caused by somehow high internal Cl^-^ ([Cl^-^]_in_) (Fig. 2B and fig. S3B). Importantly, in both cases, PSP_GABA_s were not amplified during the propagation toward both directions in dendrites (Fig. 2 and fig. S3), which was a sharp contrast to the asymmetric modulation of EPSP_glu_ (see Fig. 1). This discrepancy in the modulation of membrane depolarization spreading in a dendritic branch between glutamate- and GABA-caused ones made us to assume that the Cl^-^ conductance and/or E_Cl_ might play a critical role in processing of local depolarization in a distal dendritic branch.

**Figure 2.**
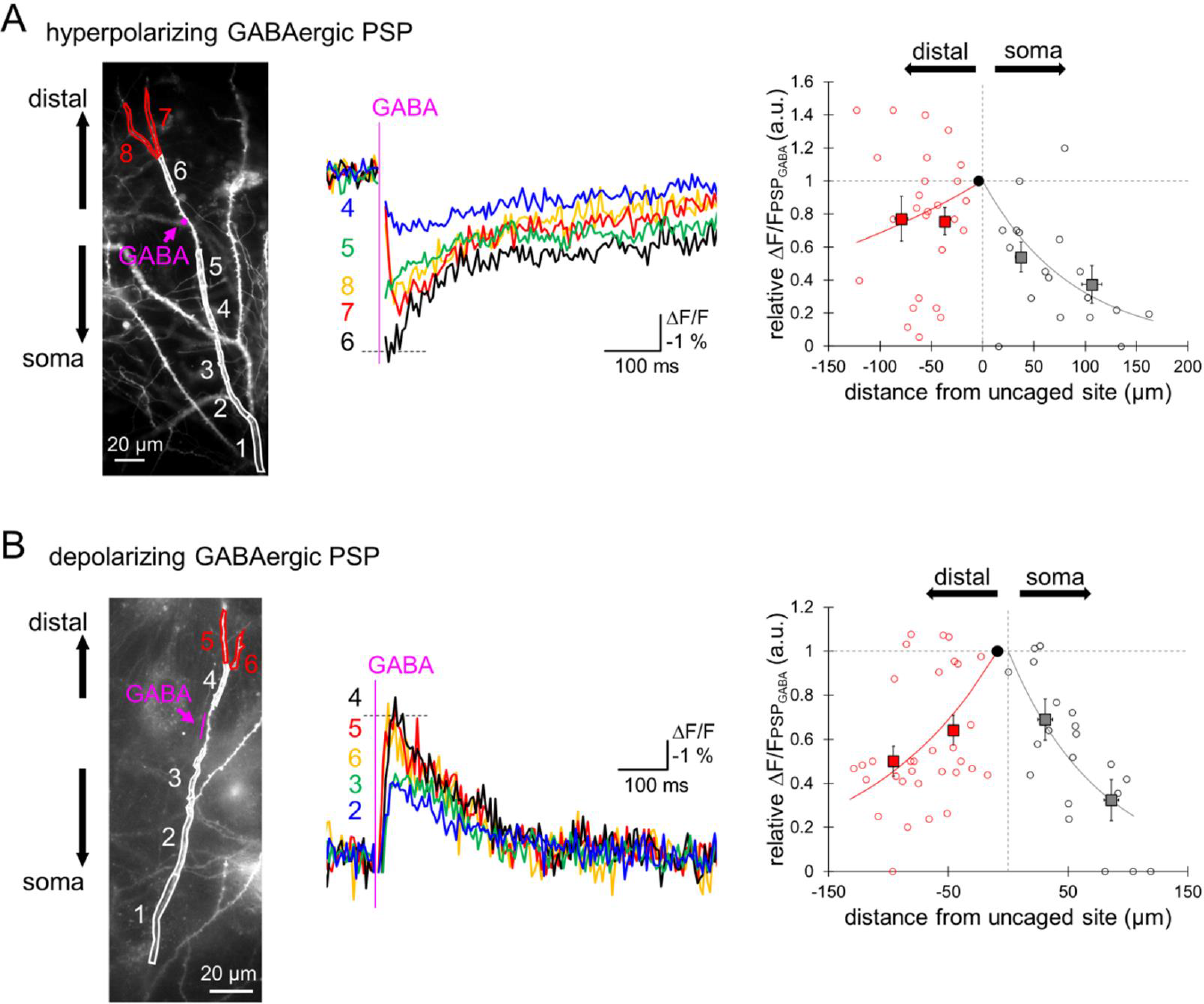
Symmetric attenuative propagation of GABAergic PSPs. **(A) and (B)** Left, dendritic segments of interest used for voltage imaging upon local GABA-evoked hyperpolarizing (A) or depolarizing (B) postsynaptic potentials (PSP_GABA_). Magenta point or line indicates the location of GABA uncaging. Middle, ΔF/F for PSP_GABA_ traces at five ROIs (4-8 for A, 2-6 for B) indicated in the left image. Right, relative ΔF/F for PSP_GABA_ is plotted against the distance from the uncaged site (6 cells, red: 25, black: 18 regions for A, 8 cells, red: 33, black: 20 regions for B). Square plots indicate averages (± SEM). See also fig. S3.

### Low [Cl^-^]_in_-dependent deeper Vm_rest_ in distal dendrites

We searched for a clue to test the idea that Cl^-^ and/or E_Cl_ affect the direction of modulation potential changes in a dendrite, first by measuring Vm_rest_ in dendrites of hippocampal neurons. For that challenging task, a direct patch-clamp recording from a very thin distal dendrite (< 1 μm in a diameter) was performed using the EGFP-fluorescence as a guide to precisely target the pipette (Fig. 3, A and B). Whole-cell current-clamp recording showed that, surprisingly, the Vm_rest_ was more negative in a dendrite (−75.5 ± 1.4 mV) than in the soma (−68.3 ± 1.4 mV, P = 0.011, Tukey test, Fig. 3C). When the dendritic Vm_rest_ was plotted against the distance from the soma, there was a tendency for Vm_rest_ to become more negative in a distal dendrite (−3.2 mV / 100 μm, Fig. 3D). In addition to the deeper Vm_rest_, leak conductance per membrane area size (measured at −70 mV under the voltage-clamp configuration) became somehow higher in a dendrite farther away from the soma (Fig. 3E), suggesting that more leak channels are basally open at the distal dendrite compared to the soma.

**Figure 3.**
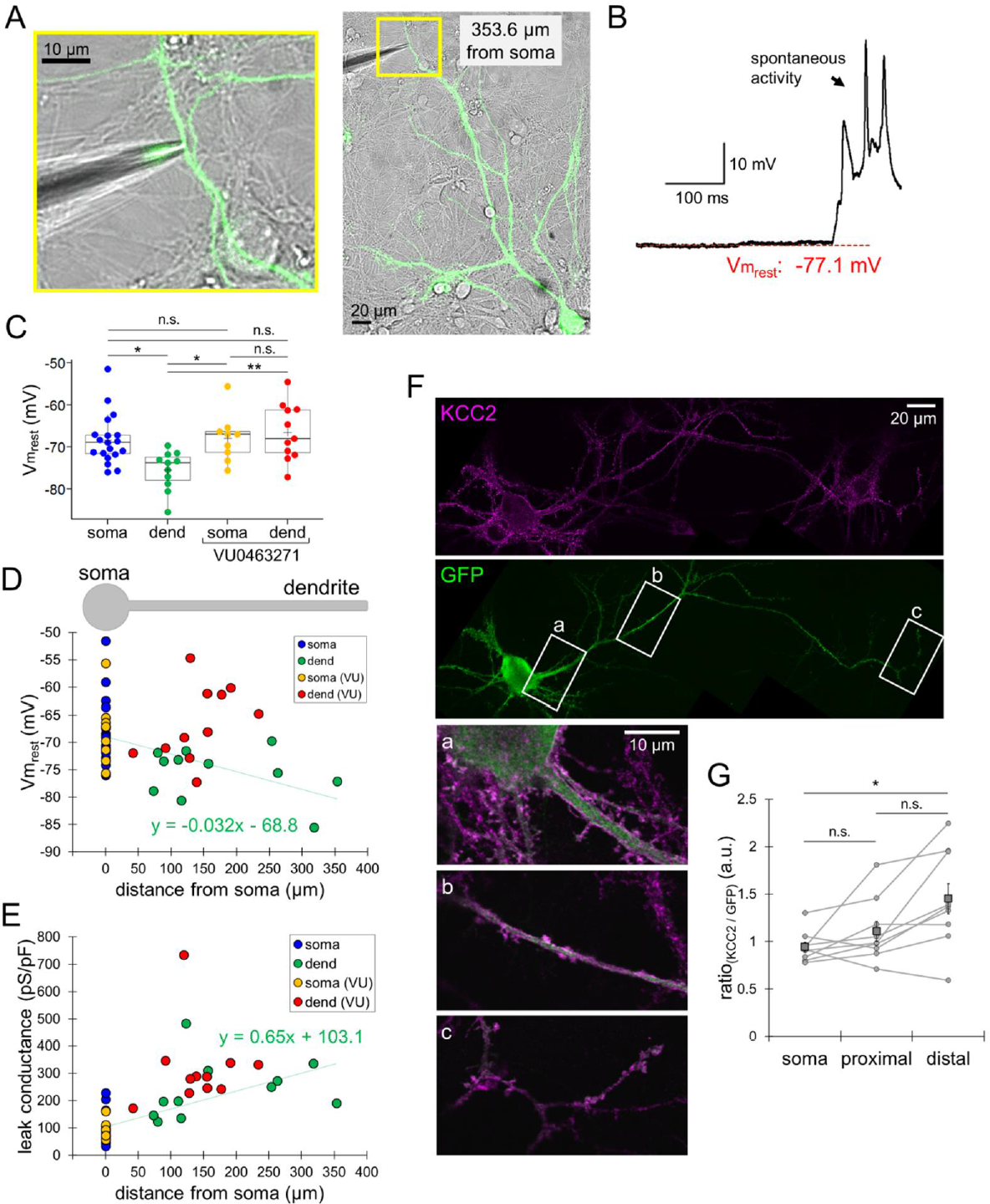
[Cl^-^]_in_-dependent more negative membrane potential in dendrites. **(A)** Image of direct patch-clamp recording from an EGFP-expressing dendrite (right) and magnified view of the patched branch in yellow (left). **(B)** V_m_ trace recorded from a dendritic branch (shown in A). **(C)** Somatic (blue, n = 19 cells) and dendritic (green, n = 11 cells) Vm_rest_ without or with 10 μM VU0463271 (soma: yellow, n = 9 cells; dendrite: red, n = 11 cells). Tukey test, *: P < 0.05, **: P < 0.01, n.s.: not significant. **(D) and (E)** Vm_rest_ (D) and leak conductance (E) plotted against the distance from soma. VU : in the presence of VU0463271. **(F)** Images of immunostained KCC2 (magenta, top) in a GFP-labeled hippocampal neuron (green, middle). Three rectangular areas (a - c) are expanded below with color-merged. **(G)** Ratio of fluorescent intensity of KCC2 to GFP in soma, proximal and distal dendrites (n = 9 cells, *: P < 0.05, n.s.: not significant, Tukey test). Square plots indicate averages (± SEM).

Taken the above results together, we hypothesized that the deeper Vm_rest_ in a distal branch might be related to the Cl^-^ conductance and/or homeostasis depending on the location of dendrite. This idea was tested using VU0463271 (10 μM), an inhibitor of KCC2, a type of Cl^-^ transporter which predominantly contributes to the low [Cl^-^]_in_ in mature neurons (19). Immunocytochemistry exhibited more abundant expression of KCC2 in distal dendrites of hippocampal pyramidal neurons than in the soma (1.5-fold, P = 0.019, Tukey test, Fig. 3, F and G). KCC2 basically establishes low [Cl^-^]_in_ by excluding Cl^-^ out of the cytoplasm, and VU0463271 is expected to bring about higher [Cl^-^]_in_. When KCC2 was inhibited, the Vm_rest_ in a dendrite became higher by ∼ 6 mV, while the somatic Vm_rest_ was not affected, resulting in almost identical potentials at dendrites and soma (dendrite: −66.5 ± 2.0 mV, soma: −67.9 ± 1.9 mV, Fig. 3, C and D). On the other hand, the tendency of higher basal leak conductance at the distal dendrite was not affected by the disruption of low [Cl^-^]_in_ (Fig. 3E). These results altogether suggested that the lower [Cl^-^]_in_ and/or higher Cl^-^ conductance in a dendrite than in the soma could be a potential cause for the deeper Vm_rest_ in dendrites.

### Deep Vm_rest_ via low [Cl^-^]_in_ and TTX-resistant Na_V_ in dendrites are critical for EPSP_glu_ amplification

Next we attempted to examine whether the deep Vm_rest_ dependent on the local Cl^-^ extrusion system in a distal dendrite is involved in the augmentation of local glutamatergic depolarizing signals. Disruption of the dendritic deeper Vm_rest_ by KCC2 inhibition using VU0463271 (10 μM) clearly, but not completely, weakened the amplification of a glutamatergic eEPSP in distal dendrites monitored by the ASAP3β imaging (ΔF/F_eEPSP_ at an uncaged site and distal site: 3.99 ± 0.39 % and 4.20 ± 0.37 %, P = 0.061, paired t-test; Fig. 4, A and D). Thus, we considered that the specific [Cl^-^]_in_ regulation and/or the resultant deeper Vm_rest_ in distal dendrites compared with the soma might play a role in the local amplification of EPSP_glu_.

**Fig. 4.**
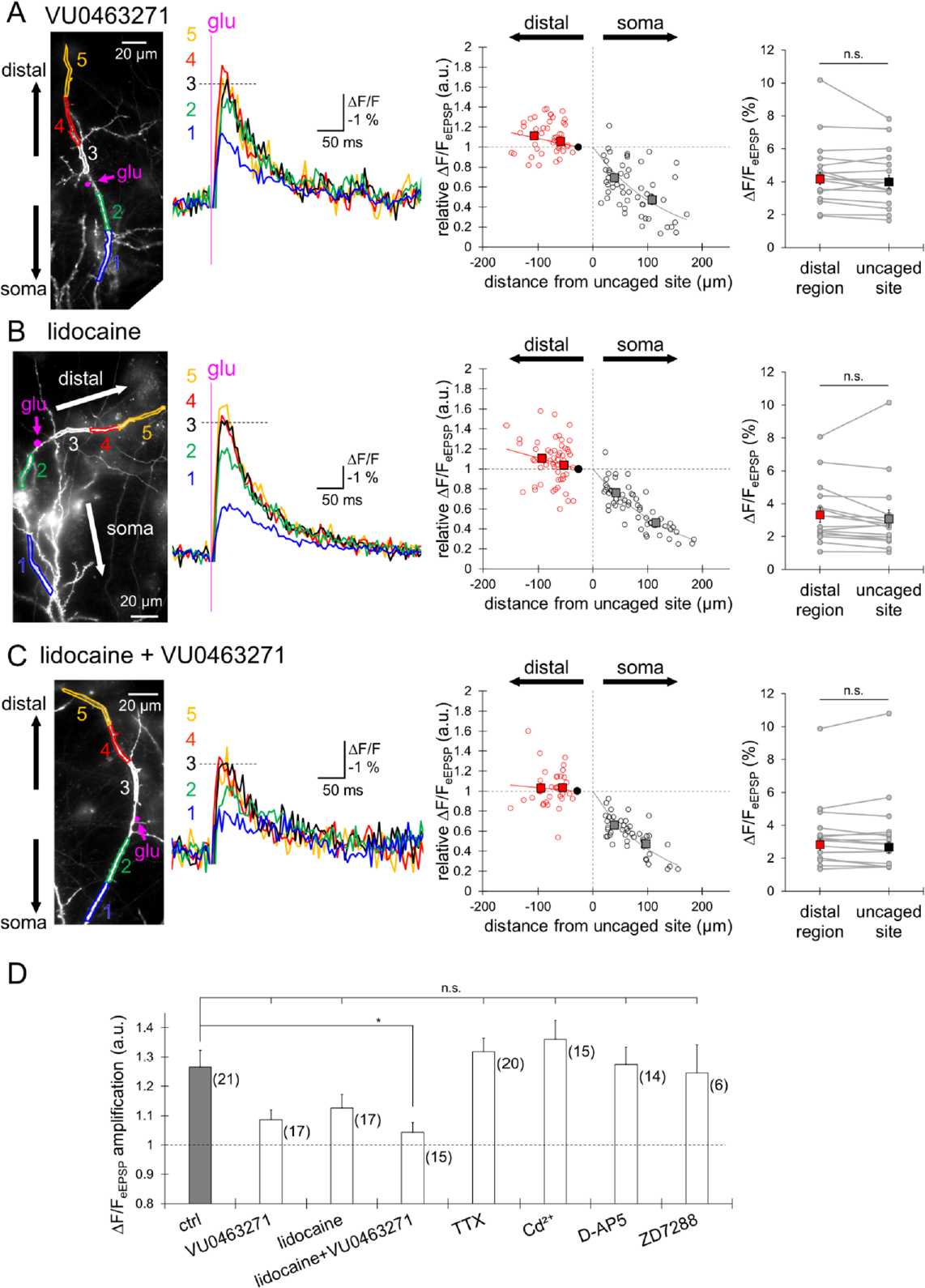
Contribution of low [Cl^-^]_in_ and TTX-resistant Na^+^ channels to EPSP amplification. **(A-C)** Left, Images for dendritic segment of interest (ROIs 1-5) and ΔF/F_eEPSP_ traces measured in the presence of VU0463271 (10 μM, A), lidocaine (5 mM, B) or both (C). Magenta points indicate the locations of glutamate uncaging. Middle, relative ΔF/F_eEPSP_ plotted against the distance from the uncaged site (17 cells, red: 44, black: 52 regions for A; 17 cells, red: 55, black: 56 regions for B; 15 cells, red: 38, black: 44 regions for C). Square plots indicate averages (± SEM). Right, absolute ΔF/F_eEPSP_ sizes in distal region of dendrites and uncaged sites (A, n = 17 cells; B, n = 17 cells; C, n = 15 cells). Paired t-test, n.s.: not significant. Square plots indicate averages (± SEM). **(D)** Statistical summary of ΔF/F_eEPSP_ amplification in distal branches with or without VU0463271 (10 μM), lidocaine (5 mM), TTX (1 μM), Cd^2+^ (100 μM), D-AP5 (100 μM), ZD7288 (100 μM). Dunnett’s test, *: P < 0.05, n.s.: not significant. Error bars are the SEM. See also fig. S4.

We next attempted to identify the remaining factor other than Cl^-^-related one, mediating the glutamate-specific dendritic augmentation of depolarization. For that aim, several ion channels were blocked by pharmacological agents. Voltage-gated Na^+^ channels (Na_V_) are known to be present in dendrites of pyramidal neurons (20), contributing to backpropagation of spikes (21). First, we applied a typical Na_V_ blocker, tetrodotoxin (TTX, 1 μM) to the external bath, and examined ΔF/F_eEPSP_ propagation by ASAP3β imaging. However, the EPSP amplification in distal dendrite was not affected by TTX (ΔF/F_eEPSP_ amplification, 1.32 ± 0.05, Fig. 4D and S4A). While TTX is a typical Na_V_ blocker used for inhibition of Na^+^ spikes, TTX-resistant Na_V_, such as Na_V_1.5, 1.8 and 1.9, are also known. Among these, at least Na_V_1.5 and Na_V_1.9 are reported to be expressed in hippocampal pyramidal neurons (22–24). In the presence of lidocaine (5 mM), which broadly inhibits Na_V_, the ΔF/F_eEPSP_ amplification in distal dendrites was not significant (P = 0.25, paired t-test, Fig. 4B), although slight augmentation was still remaining (ΔF/F_eEPSP_ amplification, 1.12 ± 0.05, Fig. 4D). Thus, TTX-resistant Na_V_ might also partially underlie the EPSP amplification in distal dendrites. Simultaneous inhibition of Na_V_ and KCC2 by lidocaine and VU0463271 almost abolished the ΔF/F_eEPSP_ amplification (ΔF/F_eEPSP_ amplification, 1.04 ± 0.03; P = 0.016, Dunnett’s test, Fig. 4C and D).

We further examined the possible involvement of other cation channels and glutamate receptors in the EPSP amplification in distal dendrites, although the EPSP size studied here was out of range for these channels. Voltage-gated Ca^2+^ channels and NMDA receptors are reported to produce dendritic Ca^2+^ and NMDA spikes upon relatively strong glutamatergic synaptic inputs, contributing to subcellular local computation by supralinear synaptic integration (6, 25). As expected, neither external administration of Cd^2+^ (Ca^2+^ channel blocker, 100 μM, Fig. 4D and S4B) nor D-AP5 (NMDA receptor antagonist, 100 μM, Fig. 4D and S4C) affected the ΔF/F_eEPSP_ amplification during propagation toward distal dendrites, suggesting that Ca^2+^ channels and NMDA receptors are not involved in the amplification of small size of EPSP here we are focusing on. In addition, inhibition of hyperpolarization-activated cation channels (HCN channels) by application of a blocker, ZD7288, failed to affect the ΔF/F_eEPSP_ amplification (Fig. 4D, and fig. S4D). Taken all these results together, it is suggested that TTX-resistant Na_V_ are critical for the EPSP amplification in distal dendrites, in cooperation with the low Cl^-^-dependent deeper Vm_rest_.

### Synergistic actions of dendritic local Cl^-^ conductances and deeper Vm_rest_ on local EPSP_glu_ amplification

Finally, we attempted to systematically understand how Cl^-^-dependent deeper Vm_rest_ and TTX-resistant Na_V_ together contribute to the local augmentation of EPSP spreading in the distal dendrite. Taking the following 2 facts into account: 1), the distal dendritic branch exhibits deeper Vm_rest_ and higher basal leak conductance (see Fig. 3); 2), the local augmentation of PSP size is specific to the depolarization mediated by Na^+^ (glutamatergic) but not by Cl^-^ (GABAergic), (see Fig. 1 and 2), we speculated that the Cl^-^ conductance at the distal dendrite might be responsible for the higher leak conductance which could bring the Vm_rest_ closer to E_Cl_. Among Cl^-^ channels, hippocampal pyramidal neurons express ClC-2 (Fig. 5A) (26), which is hyperpolarization-activated and known to regulate Cl^-^ homeostasis and membrane excitability (27, 28). Thus, as depicted in Fig. 5B, it would be possible to hypothesize that more likely activation of ClC-2 by the deeper Vm_rest_ in a distal dendrite would increase the basal Cl^-^ leak conductance, which would drive the Vm_rest_ closer to the E_Cl_ that is kept deep by the powerful local action of KCC2. To test this hypothesis mechanistically, we constructed a biophysical simple model assuming hyperpolarization-activated Cl^-^ conductance like ClC-2 and depolarization-activated Na^+^ conductance in dendritic branches (Fig. 5C; see Methods for detail of model). When an excitatory input (brief increase of conductance with a reversal potential at 0 mV) was given, an EPSP-like depolarization was caused, and importantly, the depolarization became larger during the traveling toward a region with enriched hyperpolarization-activated Cl^-^ conductances (Fig. 5, C to E). This local augmentation of depolarization in the model was suppressed either by the elimination of sensitivity of Cl^-^ conductances to hyperpolarization or by the reduction of a depolarization-caused Na_V_ activation (Fig. 5, D and E). Even when the synaptic input site is altered, the local EPSP amplification at distal branches was evident although the extent of augmentation was variable (Fig. 5E), in line with the variable size of amplification in actual data (see Fig. 1). Thus, our simple model supported the idea that the two voltage-sensitive conductances synergistically contribute to the local EPSP augmentation at distal branches.

**Figure 5.**
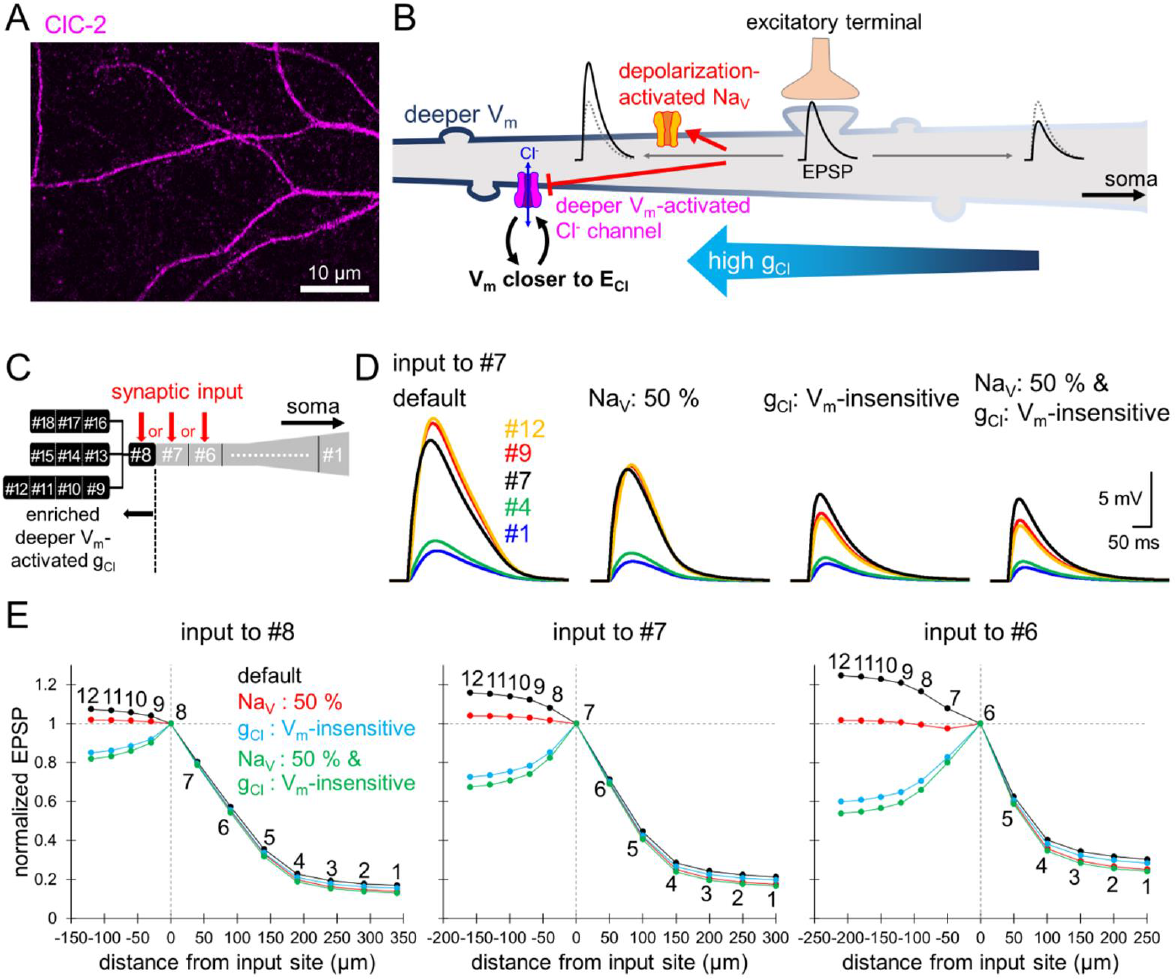
Role of Cl^-^- and Na^+^ channels in low Vm_rest_ and local EPSP amplification. **(A)** Immunostained ClC-2 image in a hippocampal neuron. **(B)** Hypothetical model for mechanisms of EPSP amplification in distal dendrites. **(C)** Geometric design of a multi-compartment model equipped with the deep V_m_-activated Cl^-^ conductance and the low-threshold Na_V_ channels. Dendrites are composed of 18 compartments, and an excitatory synaptic input was given at the compartment #6, #7, or #8. See also table S1. **(D)** Simulated EPSP traces (caused at #7) at five representative compartments (#1,4,7,9,12) indicated in C, with or without the 50 % inhibition of Na_V_ and/or the abolishment of deeper V_m_-activated Cl^-^ conductance change. **(E)** Relative amplitude of simulated EPSPs caused at #6 (right), #7 (middle), or #8 (left) plotted against the distance from the input site in 4 conditions of the model.

The above model simulation theoretically provided a prediction that the voltage-dependent fine tuning of Cl^-^ conductance in dendrites is critical for the deeper Vm_rest_ and local augmentation of EPSP_glu_. To evaluate the validity of this model prediction, we first inhibited Cl^-^ conductances by NPPB (50 μM), a blocker of relatively broad types of Cl^-^ channels including ClC-2, in addition to the Na_V_ inhibitor lidocaine, and examined by the ASAP3β imaging how the local augmentation of eEPSP in a distal dendrite is affected (Fig. 6A). Simultaneous inhibition of Na_V_ and Cl^-^ channels strongly weakened the EPSP amplification in line with the simulation results (ΔF/F_eEPSP_ amplification, 1.04 ± 0.03; P = 0.009, Dunnett’s test; Fig. 6C). Furthermore, conversely, aberrant increase of Cl^-^ conductance by applying GABA (50 μM), which would disturb the voltage-sensitive fine tuning of Cl^-^ conductance and also the Na_V_ activation through enhanced shunting effect, completely abolished the direction-dependent augmentation of EPSP_glu_ (ΔF/F_eEPSP_ amplification, 0.91 ± 0.05; P = 0.0002, Dunnett’s test; Fig. 6, B and C). Taken all these results together, our simulation of EPSP traveling in a cable equipped with 2 positive-feedback mechanisms, depolarization-activated Na^+^ conductance and hyperpolarization-activated Cl^-^ conductance, nicely predicted the cooperative action of those mechanisms to locally augment the excitatory synaptic signals, which was confirmed by the experiment.

**Figure 6.**
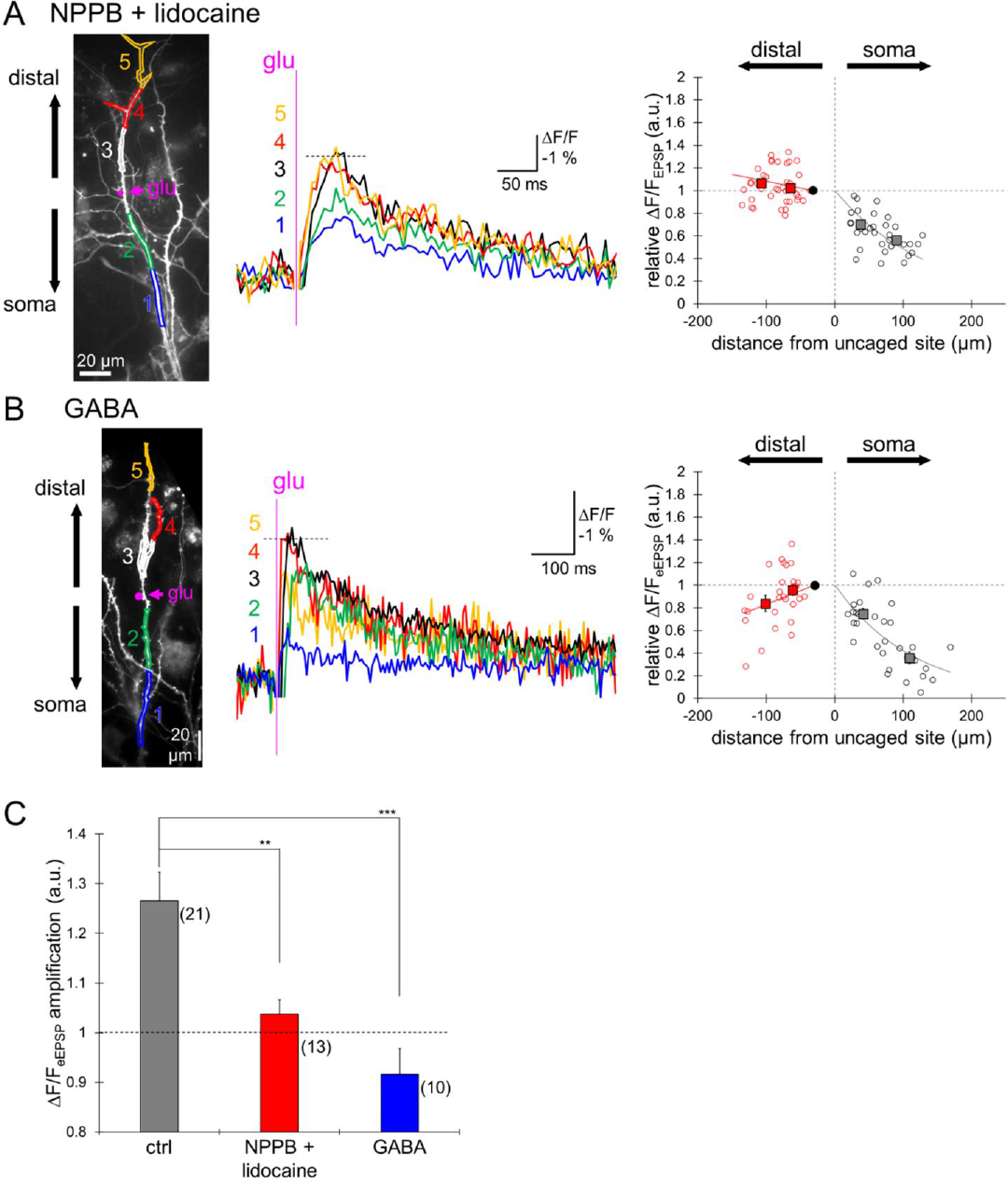
Validation of a model prediction: critical role of Cl^-^ conductance in EPSP amplification. **(A) and (B)** Left, stimulated dendritic segment in the presence of NPPB (50 μM) & lidocaine (5 mM, A), or GABA (50 μM, B). Magenta points indicate the locations of glutamate uncaging. Middle, ΔF/F_eEPSP_ traces at five ROIs (1–5) indicated in the left image. Right, relative ΔF/F_eEPSP_ plotted against the distance from the uncaged site (13 cells, red: 37, black: 34 regions for A; 10 cells, red: 29, black: 31 regions for B). Square plots indicate averages (± SEM). **(C)** Statistical summary of ΔF/F_eEPSP_ amplification in distal region with or without NPPB & lidocaine, and GABA (**: P < 0.01, ***: P < 0.001, Dunnett’s test). Error bars are the SEM.

### Direction-dependent spatio-temporal integration of synaptic inputs array

Neuronal dendrites are device for spatio-temporal integration of inputs at numerous synapses. Considering our data that glutamatergic synaptic inputs asymmetrically spread in a dendritic tree depending on the direction of propagation, we speculated that the synaptic integration might be affected by the asymmetric modulation of synaptic inputs. For example, apical dendrites of a hippocampal pyramidal neuron receive inputs from entorhinal cortex (EC) directly via perforant path (or temporo-ammonic path) at distal sites, and indirectly via the tri-synaptic circuit (mossy fibers or Schaffer collaterals) at a proximal region (29). Such a circuit organization makes a situation in which excitatory signals arrive with a spatio-temporal context: from distal to proximal. To test whether synaptic summation is affected by such a spatio-temporal context of excitatory inputs, distinct temporal orders of glutamate photolysis were performed at multiple locations along a branch (10 sites, 35 ms intervals) in the presence of TTX (1 μM). We compared the effects of three different patterns of eEPSP trains: laser spots were illuminated along a branch, 1) in a sequence from the soma toward the end (“to distal”, a to j, Fig. 7A), 2) in a sequence from the end of a branch toward the soma (“to soma”, j to a, Fig. 7A), and 3) at “random”. Fluorescent imaging with ASAP3β showed that the ΔF/F_eEPSP_ at distal region (ROI_distal_, Fig. 7A) was larger than that either at the uncaged branch or at proximal region (ROI_proximal_) in response to all three spatio-temporal patterns of stimuli (Fig. 7B). Thus, ΔF/F_eEPSP_ upon multiple stimuli on a branch is augmented during the propagation toward the distal end of dendrite, in consistent with the data for single eEPSPs. Notably, peak of ΔF/F_eEPSP_ by 10 uncagings became smaller either by lidocaine (5 mM) & VU0463271 (10 μM) or Cd^2+^ (100 μM) & D-AP5 (100 μM), compared with the control (Fig. 7C and fig. S5C), indicating several positive feedback mechanisms synergistically operate to enhance the excitatory input train during spatio-temporal summation. In addition, the location-dependent ΔF/F_eEPSP_ amplification was weakened by application of lidocaine & VU0463271 (fig. S5A), but not by Cd^2+^ & D-AP5 (fig. S5B).

**Figure 7.**
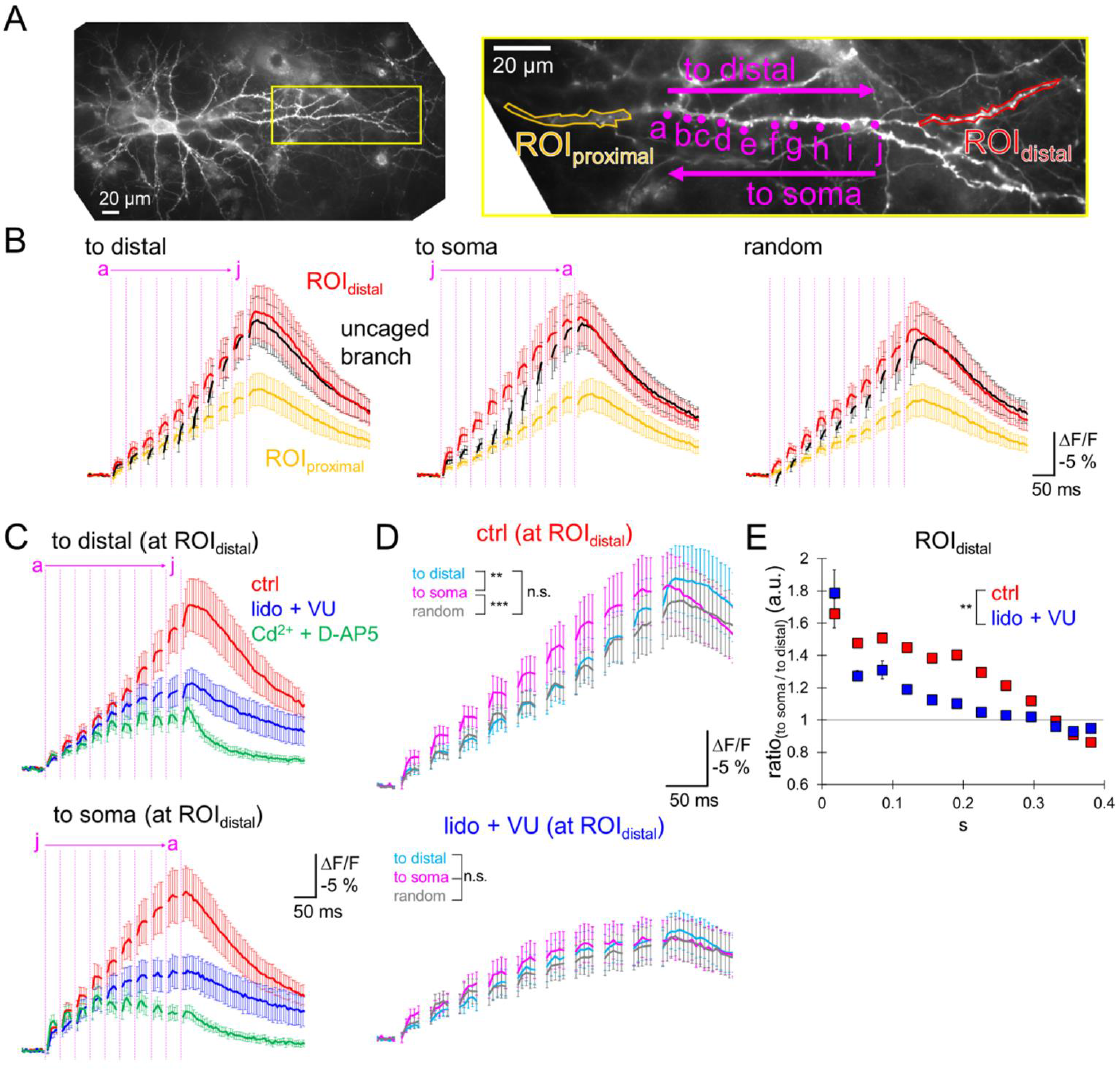
Dynamic spatio-temporal integration of synaptic inputs. **(A)** Left, image of an ASAP3β-expressing neuron, with yellow rectangular area expanded in right, showing 10 sites of local glutamate uncaging (magenta, 35 ms intervals) in three patterns: to distal (a to j), to soma (j to a) or random. **(B)** Averaged ΔF/F_eEPSP_ traces (± SEM, 17 cells) at three ROIs (uncaged branch: black, proximal: yellow, distal: red as indicated in A right) upon three patterns of glutamate uncagings. Magenta lines indicate spot laser illumination. **(C)** ΔF/F_eEPSP_ traces (mean ± SEM) at ROI_distal_ in three conditions: control (red, 17 cells), with lidocaine (5 mM) & VU0463271 (10 μM) (blue, 13 cells), and with Cd^2+^ (100 μM) & D-AP5 (100 μM) (green, 14 cells). **(D)** Comparison of synaptic integration at ROI_distal_ upon three patterns of glutamate inputs in the absence (ctrl, top) or presence of lidocaine and VU0463271 (bottom). Two-way ANOVA, **: P < 0.01, ***: P < 0.001, n.s.: not significant. **(E)** Ratio of ΔF/F_eEPSP_ signals (mean ± SEM) upon opposite directions of glutamate uncagings (“to soma” relative to “to distal”) at ROI_distal_ in the absence (ctrl) or presence of lidocaine & VU0463271. Two-way ANOVA, **: P < 0.01. See also fig. S5.

Furthermore, pyramidal neuronal dendrites exhibited distinct augmentation of EPSP summation depending on the spatio-temporal order of synaptic activation. Although the ΔF/F_eEPSP_ at ROI_distal_ reached similar peaks in two conditions, “to soma” and “to distal” (Fig. 7D and fig. S5D, P > 0.05, paired t-test), the signal at a distal branch (ROI_distal_) steeply summated during glutamate uncagings in a sequence of “to soma” compared to “to distal” (Fig. 7D, P < 0.01, two-way ANOVA), but not at a proximal branch (ROI_proximal_, fig. S5E). The ratio of the eEPSP train trace (at ROI_distal_) for “to soma” divided by that for “to distal”, reflecting the preference of “to soma” direction for effective summation of excitatory inputs, was kept higher than 1 during the EPSP array (Fig. 7E). Importantly, such a context-dependent augmented summation was clearly weakened by lidocaine & VU0463271 (Fig. 7D, P > 0.05, two-way ANOVA; and Fig. 7E, P < 0.01, two-way ANOVA). Thus, the Na_V_- and low [Cl^-^]_in_-dependent ΔF/F_eEPSP_ amplification seems to make it possible for a distal branch to decode the spatio-temporal context of synaptic inputs, resulting in local computation in a distinct manner.

## Discussion

In this study, using voltage imaging with a genetically-encoded probe ASAP3β combined with local glutamate or GABA uncaging, subcellular patch-clamp recordings, and biophysical model simulation, we explored how subthreshold size of glutamatergic and GABAergic synaptic inputs propagated in hippocampal neuronal dendrites. Imaging analysis of membrane potential changes at multiple dendritic branches demonstrated direction-dependent asymmetric propagation of excitatory synaptic inputs in a dendrite, that is, EPSPs attenuated during propagation toward the soma, while amplified toward the distal end. Moreover, the EPSP amplification depended on synergistic actions of TTX-resistant Na_V_ and low [Cl^-^]_in_-related Cl^-^ conductance. Thus, the data presented in this study showed asymmetric modulation of glutamatergic synaptic inputs in contrast to the conventional view established from the classical cable theory: subthreshold membrane potential changes gradually attenuate during spreading in dendrites. Together, our findings provide a novel concept of local modulation of synaptic integration relying on a unique Cl^-^-dependent mechanism in a single neuron.

To examine the spatio-temporal propagation pattern of synaptic inputs, we used fluorescent voltage imaging with a GEVI. Patch-clamp electrophysiology has been a powerful technique to precisely record electrical signals at a given position in a neuron, although it requires a sophisticated technique. On the other hand, GEVIs enable us to measure membrane potential changes much more easily even from multiple subcellular regions at the same time, that is very difficult to perform by conventional patch-clamp recordings. In this decade, several types of GEVI, including ASAP (13), have been developed, such as ArcLight (30), Ace2N-mNeon (14), Voltron (31), and so on (15, 32). Taking advantage of this merit, recent studies applied GEVIs to various preparations *in vivo* and *in vitro,* demonstrating neuronal excitability changes dependent on behavioral states in animals (32). Moreover, GEVIs could be non-invasive compared with whole-cell patch-clamp recordings which is typically accompanied with washout of cytoplasmic components. As demonstrated in this study, [Cl^-^]_in_ is critical for the local EPSP amplification (see Fig. 3-7), and we speculate that monitoring of membrane potential without intracellular dialysis must have been essential for the detection of such cellular functions. In addition, the EPSP amplification demonstrated here operates even for small EPSPs (locally 2 ∼ 5 mV and hence ∼ 1 mV or smaller at the soma) in a very local branch of distal dendrites specifically, which cannot be recognized by the somatic recordings. Therefore, the lack of useful technique like voltage imaging based on GEVIs must have veiled the Cl^-^-dependent asymmetric EPSP amplification at a distal dendrite.

Previous studies have suggested local augmentation of synaptic inputs by active conductances via voltage-gated cation channels such as Na^+^ and/or Ca^2+^ channels and NMDA receptors (4, 6, 25, 33). Notably, such augmentation of local excitatory inputs emerges when the input size is relatively large (∼ 5 mV or larger when recorded at the soma and hence ∼ 20 mV at the stimulated site). On the other hand, as shown in this study, eEPSPs caused here by spot laser illumination on a spine, was not so large in amplitude: ∼ 3 % ΔF/F in average, corresponding to ∼ 5 mV depolarization at the stimulated site, and the local EPSP amplification relied on the TTX-resistant Na^+^ channels. Interestingly, among three subtypes of such Na^+^ channels (Na_V_1.5, 1.8 and 1.9), Na_V_1.9 is especially activated at more negative membrane potential (−70 ∼ −60 mV) (34, 35). Taking it into consideration that activations of typical Na_V_ and Ca^2+^ channels are triggered at −40 ∼ −50 mV, these channels are unlikely to be involved in the amplification of small size of EPSPs. Thus, distinct subtypes of Na_V_ with different activation profile might provide neurons wide range of locally computation.

In contrast to the conventional view of dendritic computation focusing on cation channels, this study identified a unique mechanism of local augmentation of excitatory inputs in a distal dendrite: low [Cl^-^]_in_- and Cl^-^ conductance-dependent enhancement of depolarization in distal dendrites (see Fig. 4-6). Our biophysical model simulation indicated that a type of hyperpolarization-activated Cl^-^ channel, ClC-2, could constitute a kind of positive-feedback loop to stably keep the membrane potential negative as close to the E_Cl_. Indeed, direct subcellular patch-clamp recordings from dendritic branches demonstrated the deeper Vm_rest_ dependent on low [Cl^-^]_in_ and higher leak conductances at distal branches in a distance-dependent manner (see Fig. 3). Such a positive-feedback hyperpolarizing system is switched-off upon a depolarization, resulting in indirect enhancement of depolarizing responses (see Fig. 5). The specific subtype of Cl^-^ channels in the asymmetric EPSP modulation, here we could not determine because of the lack of specific blockers, is an important issue to be addressed in a future study.

In addition to the involvement of Cl^-^ conductance, E_Cl_ may also be altered at distal dendrites contributing to deeper Vm_rest_ there because of the lower [Cl^-^]_in_ which is established by KCC2 activity, as reported previously (36–39). More abundant expression of KCC2 in spines and vicinity of spines (37, 40) and/or more efficient Cl^-^ extrusion by KCC2 might operate at distal dendrites based on the distinct surface/volume ratio, as shown by Báldi et al. (41). Furthermore, regulation of KCC2 by phosphorylation and dephosphorylation in distal dendrites (36, 42, 43) may also give rise to diverse modes of Cl^-^ homeostasis, potentially impacting the local processing of synaptic potentials in a branch. As shown in Fig. 1 and 2, in contrast to the asymmetric modulation of EPSP_glu_s, depolarizing PSP_GABA_s symmetrically attenuated during propagation in dendrites. Depolarizing PSP_GABA_s would have been caused by unusually high [Cl^-^]_in_, in a manner similar to the situation with VU0463271 (see Fig. 3), so that the E_Cl_ should be higher and then the deep Vm_rest_ at a distal dendrite would also be disturbed, precluding one of the two critical positive-feedback mechanisms for the local amplification of depolarization.

As demonstrated in this study (see Fig. 7), hippocampal and cortical pyramidal neurons exhibit supralinear synaptic integration upon multiple inputs in local dendrites dependent on active conductances (5, 6), whereas other types such as dentate gyrus granule cells and cerebellar interneurons show linear or sublinear summation (44–46). Accordingly, among hippocampal culture preparation consisting of several types of neurons, pyramidal cells, but not apparent granule cells, selectively exhibited the local asymmetric EPSP modulation (see Fig. 1). Thus, distinct neuronal types seem to be equipped with specific machinery for local dendritic computation. Asymmetric propagation of EPSPs may play a role in associative induction of synaptic plasticity such as long-term potentiation or depression. EPSPs caused at a dendrite sequentially in the direction toward the soma, likely cause an AP firing in cortical pyramidal neurons (47). Furthermore, the long-term plasticity at excitatory synapses tends to be induced by dendritic spike firing in a distal thin branch caused by clustered inputs, rather than distributed ones (48, 49). Thus, asymmetric modulation of EPSPs revealed here might associate a given pattern of EPSPs arriving at a distal dendrite, contributing to the local dendritic spikes and long-term plasticity.

## Methods

### Plasmid construction

Plasmids were constructed by standard molecular biology techniques. The coding DNA for ASAP1 was obtained from Addgene (#52519), and some modifications for higher voltage sensitivity (D145Δ, L146G, S147T, N149R, S150G, and H151D) in an improved version ASAP3 (18) were introduced by PCR-mediated mutations. Modified-ASAP (ASAP3β) or original ASAP1 cDNAs was inserted into the vector made from pAAV-MSC by replacement of CMV promotor and beta-globin intron region by CA promotor and intron region from pCAGGS, followed by insertion of WPRE sequence before hGH pA region.

### HEK293T cells

HEK293T cells were cultured at 37 °C in humidified air containing 5 % CO_2_ in DMEM (Nakalai Tesque, Kyoto, Japan) supplemented with 10 % (v/v) FBS (NICHIREI BIOSCIENCES, Tokyo, Japan) and penicillin (100 U/mL)-streptomycin (0.1 mg/mL, Nakalai Tesque). For testing voltage-dependent fluorescence changes, HEK293T cells were transfected with original ASAP1 or modified-ASAP (ASAP3β) using Lipofectamine 3000 (1 μg DNA, 2 μL P3000 reagent, 1.5 μL Lipofectamine for 35 mm dish; Thermo Fisher Scientific, Waltham, MA, USA).

### Hippocampal neuron culture and transfection

The animals were maintained and treated according to the NIH guide for the care and use of laboratory animals, and to the ethical guidelines on animal experimentation of Kyoto University. Hippocampi were dissected out from male and female newborn Wistar rats and incubated in Ca^2+^ and Mg^2+^-free Hank’s balanced salt solution (CMF-HBSS) containing 0.1 % trypsin for 5 min at 37 °C. Neurons were dissociated by trituration with a fire-polished Pasteur pipette and seeded on poly-D-lysine (Sigma-Aldrich, St. Louis, MO, USA)-coated glasses in Neurobasal plus medium containing 2 % B27 plus supplement (Thermo Fisher Scientific) and penicillin (100 U/mL)-streptomycin (0.1 mg/mL, Nakalai Tesque). Some cells were transfected with plasmids by means of electroporation with NEPA21 (Nepa Gene, Chiba, Japan). 1 × 10^6^ cells were mixed with plasmids (6 μg ASAP3β plasmid or 3 μg pCAsalEGFP (50)) in 100 μL CMF-HBSS. Square electric pulses were applied at 300 V (pulse length, 0.5 ms; two pulses; interval, 50 ms; voltage decay rate, 10 %), followed by additional pulses at 20 V (pulse length, 50 ms; five pulses; interval, 50 ms; voltage decay rate, 40 %; polarity exchanged pulse). Transfected cells were then cultured on the glasses together with the non-treated ones. The cells were cultured at 37 °C in humidified air containing 5 % CO_2_. One day after seeding, 75 % of the medium was replaced with fresh one. Since then, half of the medium was replaced every 4 days.

### Electrophysiology

Whole-cell voltage-clamp recording from an ASAP-expressing HEK293T cell was performed with an amplifier (EPC10; HEKA Elektronik GmbH, Reutlingen, Germany) in an extracellular solution containing (in mM) 145 NaCl, 5 KOH, 2 CaCl_2_, 1 MgCl_2_, 10 Hepes and 10 glucose (pH 7.3) at room temperature (20-24 °C). Patch pipettes (3-5 MΩ) were filled with an internal solution containing (in mM) 147 CsCl, 5 EGTA, 10 Hepes, 15 CsOH, 2 ATP, 0.2 GTP (pH 7.3). The membrane potential (a holding potential of - 70 mV) was changed by step voltage depolarizations and hyperpolarizations to a voltage (−100 mV to 50 mV) for 30 ms.

In recordings from hippocampal neurons in culture for 2-3 weeks, we used an extracellular solution containing (in mM) 120 NaCl, 5 KOH, 2 CaCl_2_, 1 MgCl_2_, 10 Hepes and 10 glucose (pH 7.3). Patch pipettes (3-5 MΩ, somatic recordings; 20-25 MΩ, dendritic recordings) were filled with an internal solution containing (in mM) 120 K-gluconate, 7 KCl, 5 EGTA, 10 Hepes, 2 ATP, 0.2 GTP (pH 7.3 adjusted with KOH). In some experiments, TTX (1 μM, FUJIFILM Wako Pure Chemical Corporation, Tokyo, Japan), lidocaine (5 mM, Nakalai Tesque), D-AP5 (100 μM, Tocris Bioscience, Bio-Techne SRL, Minneapolis, MN, USA), Cd^2+^ (100 μM, FUJIFILM Wako Pure Chemical Corporation), ZD7288 (100 μM, Tocris bioscience), VU0463271 (10 μM, Tocris Bioscience) and NPPB (50 μM, Tocris Bioscience) were used to inhibit Na^+^ channels, NMDA receptors, Ca^2+^ channels, HCN channels, IRK channels, KCC2 and Cl^-^ channels, respectively. GABA (50 μM, Nakalai Tesque) was used to activate GABA_A_ receptors.

### Fluorescent imaging

Fluorescence imaging was performed using an inverted microscope IX71 (Olympus, Tokyo, Japan) through a 40× 0.95-NA objective (Olympus) for HEK293T cells, or a 60× 1.42-NA oil-immersion objective (Olympus) for hippocampal neurons. Fluorescent excitation was delivered using a white LED (SOLA; Lumencor, OR, USA) with 15 mW/mm^2^ through a 450-480 nm filter. Fluorescent images were obtained with a Zyla4.2 sCMOS camera (Andor Technology Ltd, Belfast, UK) at 1500 Hz (128 × 128 pixels, binning 2 × 2) for HEK293T cells, at 400 Hz (512 × 512 pixels, binning 2 × 2) or at 200 Hz (1392 × 1040 pixels, binning 2 × 2) for hippocampal neurons and analyzed with SOLIS (Andor) or Image J software (NIH, Bethesda, MD, USA).

The LED illumination in our self-customized fluorescent imaging system is designed to be assembled around the center of view field so that high fluorescent signal is attained as possible even for a very fast image acquisition. As a result, the absolute fluorescent signals around the edge of view field tended to be smaller, potentially giving rise to a tendency for the signal to noise ratio to become lower at those dimmer regions. However, we confirmed that, as shown in fig. S2, spatial pattern of ΔF/F_eEPSP_ signals in dendrites was little affected by the relative brightness of fluorescence altered by moving the microscope.

### Glutamate or GABA uncaging

For glutamate or GABA uncaging, MNI-caged-L-glutamate (300 μM, Tocris Bioscience) or DPNI-caged-GABA (200-500 μM, Tocris Bioscience) was applied to the external solution. Uncaging was caused by 405 nm laser illumination (Single line laser; Rapp OptoElectronic GmbH, Wedel, Germany) with 4-6 mW laser power (51, 52). Glutamate uncaging was performed by laser illumination at a 2 μm diameter spot (0.2 ms duration; or 1 ms in the presence of external GABA) around 200 μm from the soma, while GABA was uncaged by a spot or a line scan (0.5-5 ms duration). The location of laser illumination was controlled by a localized photomanipulation system based on Galvano mirrors (UGA-42 Firefly; Rapp OptoElectronic) implemented in SysCon software (Rapp OptoElectronic).

### Immunocytochemistry

Cultured neurons were fixed with 4 % paraformaldehyde in phosphate buffered saline, permeabilized with 0.5 % Tween 20, then blocked with 1 % goat serum, and finally labeled with primary and secondary antibodies. The following antibodies were used: rabbit polyclonal antibodies (pAb) against KCC2 (1:1000, Sigma-Aldrich), ClC-2 (1:5000, Alomone Labs, Jerusalem, Israel), chicken pAb against GFP (1:2000, Sigma-Aldrich), Alexa 568-conjugated pAb against rabbit IgG (1:400, Thermo Fisher Scientific) and Alexa 488-conjugated pAb against chicken IgG (1:400, Thermo Fisher Scientific). Fluorescent images were recorded with a confocal laser microscope (FV1000 imaging system, Olympus, Japan), and analyzed using ImageJ software (NIH). The neuronal dendrites were defined as the area positive for GFP.

### Simulation of simple dendrite model with voltage-dependent Na^+^ and Cl^-^ channels

Simple model of a dendrite equipped with the depolarization-activated Na_V_ and the hyperpolarization-activated Cl^-^ channels was constructed and simulated using the *NEURON* simulator (53) version 8.2 operating under the Python programming language. Here we assumed a multi-compartment model consisting of a soma and 18 dendritic sections with single branch point where 3 daughter branches stemmed out (see Fig. 5C), one of which had 120 μm and the other two 90 μm in length, yielding total arborization length of 500 or 470 μm long. The diameter of a dendritic branch was assumed to be 2.5 μm at the most proximal one, narrowing down towards the distal end until it reaches 0.5 μm at 200 μm away from the soma. Cytosolic resistivity was here set at 250 Ω·cm, and membrane capacitance 0.8 mF/cm^2^. Membrane of individual compartments were equipped with TTX-resistant voltage-gated Na^+^ channel Na_V_1.9, hyperpolarization-activated Cl^-^ channel like ClC-2, Na^+^/K^+^ pump, KCC2, and linear leak conductances for Na^+^, K^+^, and Cl^-^ (for detailed parameters of each components, see table S1).

Individual components in each compartment at the membrane potential (V_m_) were expressed in the model based on the equilibrium potentials for Na^+^ (E_Na_), K^+^ (E_K_), and Cl^-^ (E_Cl_) by implementing the equations which were already published in previous studies, as described below.

For TTX-resistant persistent Na channels, Na_V_1.9 model (54) was here used:

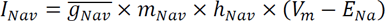

, where 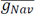, m_Nav_, h_Nav_ are the maximal conductance, voltage-dependent activation and inactivation probability factors, respectively. Voltage-dependent changes of m_Nav_ and h_Nav_ were defined by the rate constants (α and β) as described by the following equations:

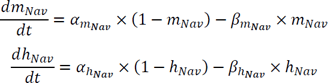

, where forward (α) and reverse (β) rate constants for m_Nav_ and h_Nav_ were assumed as below:

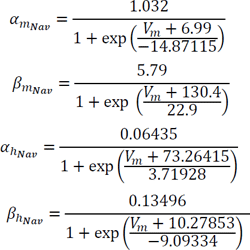

For hyperpolarization-activated Cl^-^ component, a modified biophysical model for ClC-2 (55) was incorporated:

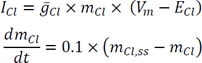

, where 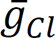 is the maximal conductance, m_Cl_ is the relative open probability factor of the channel, and its steady state at V_m_, m_Cl,ss_ is defined as:

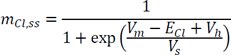

, where V_h_ and V_s_ are parameters to define the voltage-sensitivity of the channel activation.

For KCC2, K^+^ and Cl^-^ fluxes were calculated as a product of the parameter for its anti-porting efficiency (U_KCC2_) and the difference between their equilibrium potentials as demonstrated in Gentiletti et al. (56):

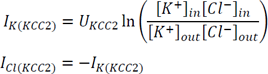

Na^+^/K^+^ pump activity was calculated based on the formulations in Takeuchi et al. (57), as listed below,

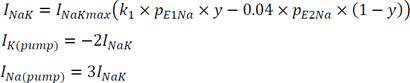

State transitions of the pump were defined by rate constants as below:

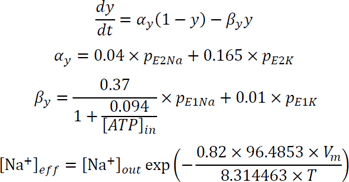

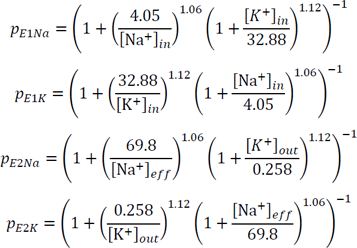

In our model, calculation of the Na^+^/K^+^ pump activity was specifically implemented in Neuron simulator, by modifying the program code presented in Botta et al. (58).

Notably, here we assumed the hyperpolarization-activated Cl^-^ channels only at distal compartments (see Fig. 5C). An EPSP was represented as a double exponential conductance change (reversal potential at 0 mV) at a single compartment (#6-8 in Fig. 5C), defined by the rise and decay time constants of 10 and 80 ms, respectively, and the peak conductance was optimized to cause similar size of EPSP (∼ 12 mV) at the stimulated compartment. The model was simulated in *Neuron* platform with backward Euler method (dt = 0.025 ms). In some simulations to evaluate the role of the voltage-dependent Na^+^ and Cl^-^ conductances in the model, the maximal conductance of Na_V_ was halved and/or the Cl^-^ conductances at distal branches were set constant.

### Data analysis of ASAP fluorescence change

Dendritic regions were selected for analysis as ROIs in ImageJ, and F_dend_ was calculated as the mean intensity of fluorescence in each ROI which was subtracted by background noise defined as the mean of fluorescence intensity at the area nearby each ROI. During image acquisition of ASAP-expressing cells, fluorescence gradually decreased due to photobleaching, which was defined as F_base_ obtained by double exponential curve fitting of the total fluorescence change as follows:

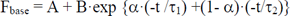

All above parameters (A, B, α, τ_1_ and τ_2_) were determined by the least square method from the time course of fluorescence change before the uncaging. The relative change of fluorescence (ΔF/F, %) was

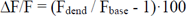

In this study, relative fluorescence change of ASAP is represented as −ΔF/F because of the negative relation of fluorescence against membrane potential change. −ΔF/F traces in individual ROIs were obtained by averages of 3-7 trials. Adjacent area to the laser spot was excluded from analysis because of an artifact by laser illumination. Distance between two sites of interest was measured using ImageJ (NIH). Data fitting for ΔF/F against the distance from the uncaged site was performed by Igor Pro8 (WaveMetrics, Lake Oswego, OR, USA) or Microsoft Excel.

### Statistics analysis

Statistical significance of differences between groups was tested by two-tailed paired Student’s t-test, Dunnett’s test, Tukey test or two-way ANOVA, and P < 0.05 was considered as significant. Throughout the project, data are presented as mean ± standard error of the mean (SEM) unless otherwise stated. In all figures, symbols with error bars indicate mean ± SEM; asterisk, P < 0.05; double asterisk, P < 0.01; triple asterisk, P < 0.001. These analyses were performed by Microsoft Excel and Igor Pro8 (WaveMetrics).

## Acknowledgments

We thank to Drs. M. Midorikawa and T. Inoshita for critical reading of the manuscript and helpful comments.

## Author contributions

MM performed most imaging and patch-clamp experiments and data analysis; RH performed model simulation and analysis; SK conceptualized and administrated the study; SK and MM interpreted data and wrote the manuscript. All authors approved the final version.

## Data and materials availability

All plasmids used in this study except for those available at Addgene can be requested to the corresponding author. All data and any additional information required to reanalyze the data reported in this paper will be shared by the corresponding author upon request. Program codes for simulating the simple biophysical multi-compartment model will be available at the website of the corresponding author.

