## Supplementary figures and table for "Cl^-^-dependent amplification of excitatory synaptic potentials at distal dendrites revealed by voltage imaging"

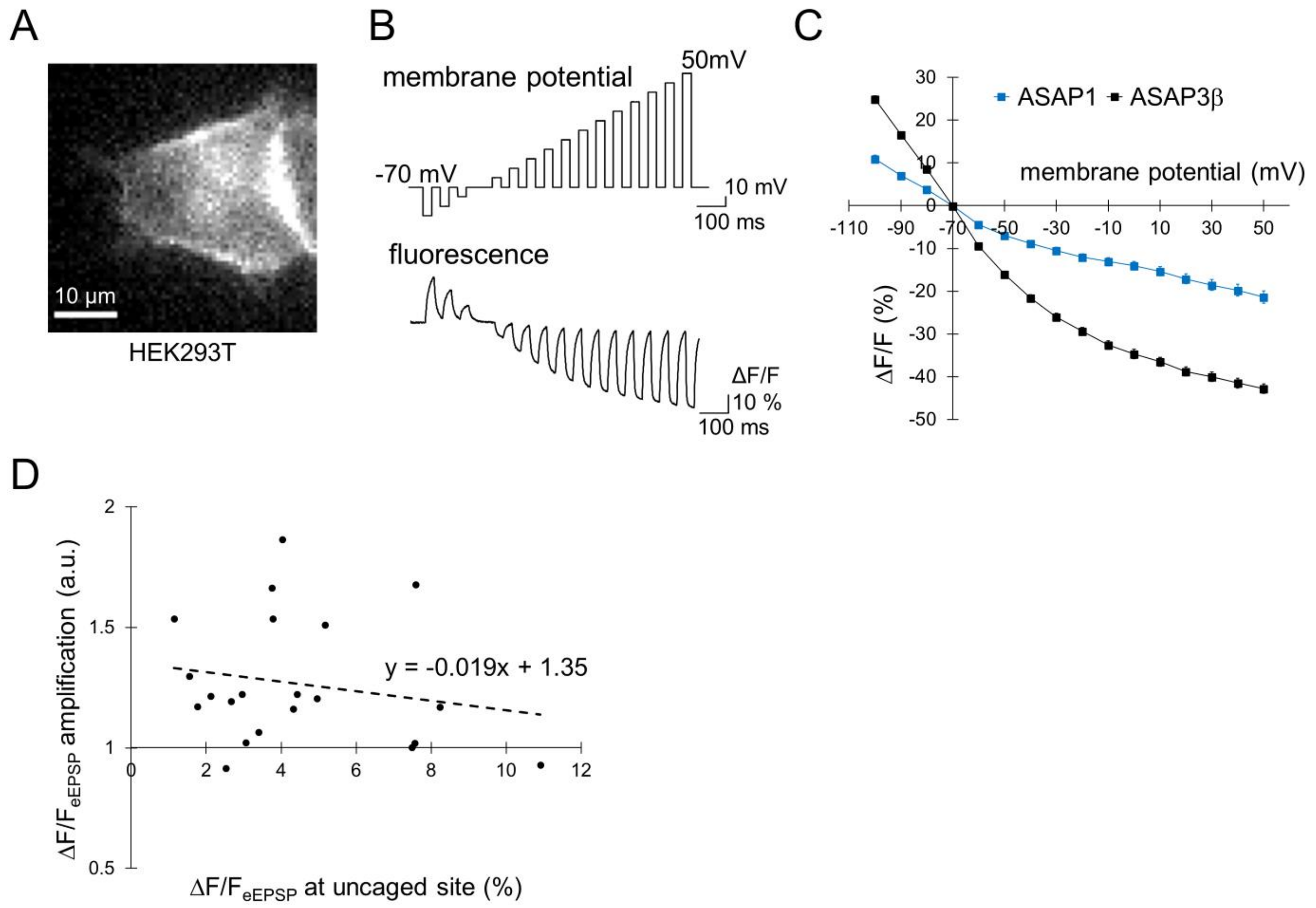


**Fig. S1. Voltage sensitivity of ASAP3β and ΔF/$F_{eEPSP}$ amplification independent of the eEPSP amplitude**

**(A)** Fluorescent image of a HEK293T expressing ASAP3β.

**(B)** Fluorescent intensity changes of ASAP3β (bottom) in a whole-cell voltage clamped HEK293T cell in response to voltage steps to -100 ~ 50 mV from -70 mV (top). Fluorescent changes (ΔF) are normalized to the fluorescence intensity at the -70 mV holding potential (F).

**(C)** Relative fluorescence changes (ΔF/F) of ASAP1 and improved version (ASAP3β) are plotted against membrane potentials (ASAP1, n = 5 cells; ASAP3β, n = 6 cells). Error bars are the SEM.

**(D)** Relative ΔF/$F_{eEPSP}$ amplification in a distal neuronal dendrite plotted against absolute ΔF/$F_{eEPSP}$ around the uncaged site (n = 21 cells).

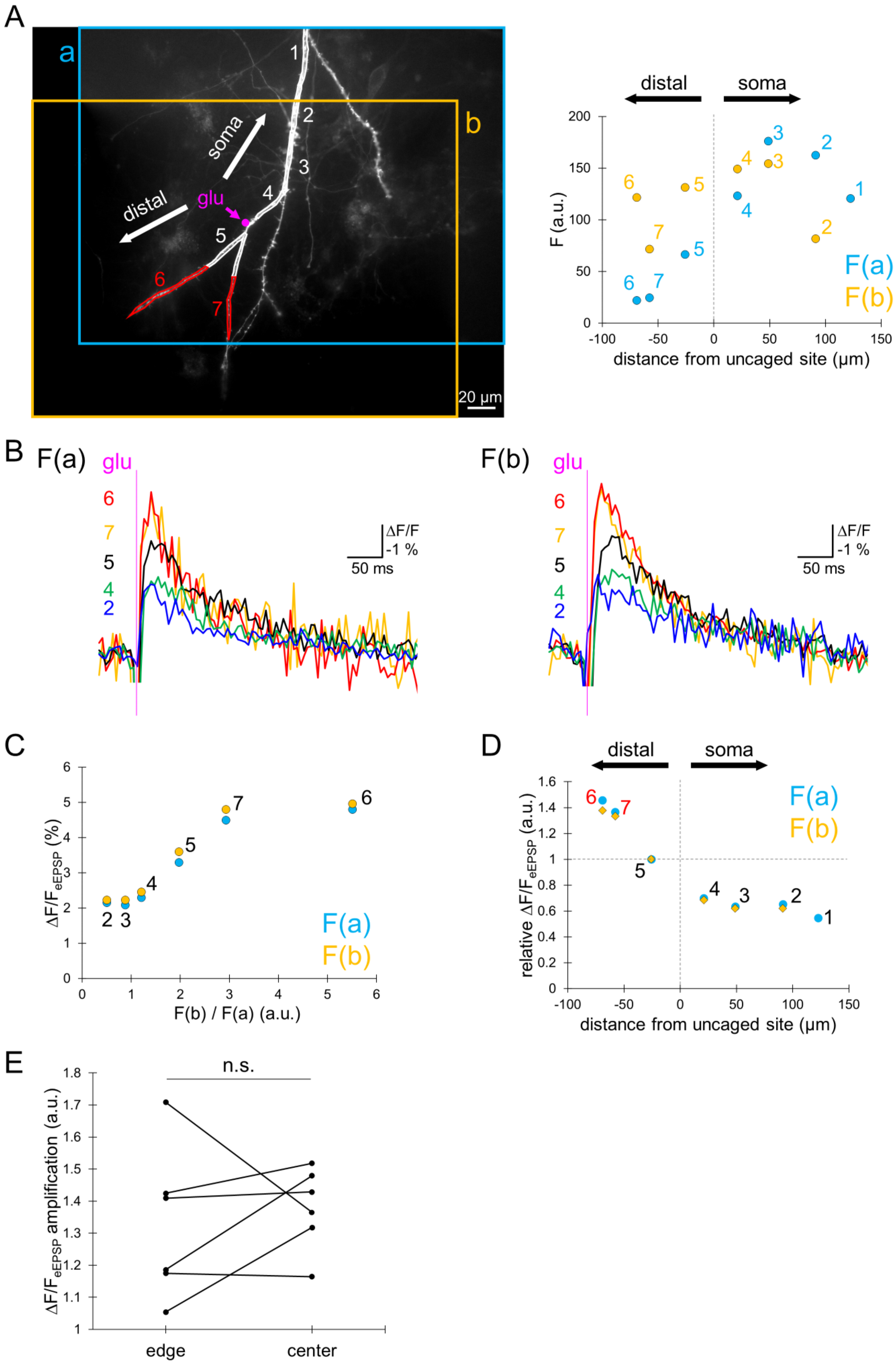
A
a
b
soma
distal
glu
1
2
3
4
5
6
7
20 μm
distal
soma
F (a.u.)
distance from uncaged site (μm)
F(a)
F(b)
B
F(a)
F(b)
glu
ΔF/F
-1 %
50 ms
C
ΔF/F_eEPSP (%)
F(b) / F(a) (a.u.)
D
relative ΔF/F_eEPSP (a.u.)
E
n.s.
ΔF/F_eEPSP amplification (a.u.)
edge
center

**Fig. S2. Voltage imaging quantification unbiased by image acquisition setting**

**(A)** Left, image acquisitions of a branch with different view fields (a: sky blue and b: orange square). Magenta point indicates the location of glutamate uncaging. Right, fluorescence intensity at each ROI showing substantial difference depending on the relative location of view field.
**(B)** $\Delta F/F_{eEPSP}$ traces at five ROIs (2,4,5,6,7 shown in A), in two distinct image acquisition settings, (a) and (b).
**(C)** $\Delta F/F_{eEPSP}$ amplitude at each ROI plotted against the ratio of absolute fluorescent intensities (F(a) to F(b)) at different image acquisitions.
**(D)** Relative $\Delta F/F_{eEPSP}$ plotted against the distance from the uncaged site. Sky blue points are data from data (a), and orange points from data (b). Responses at individual ROIs are normalized to the $\Delta F/F_{eEPSP}$ peak at ROI 5.
**(E)** Summary of the $\Delta F/F_{eEPSP}$ amplification in a distal branch between two distinct image acquisition settings. "edge" represents the image acquisition setting in which ROIs at distal branch from the uncaged site are located close to edge of image, and "center" represents that ROIs in distal branch are located closer to the center of image (n = 6 cells; n.s.: not significant, paired t-test).

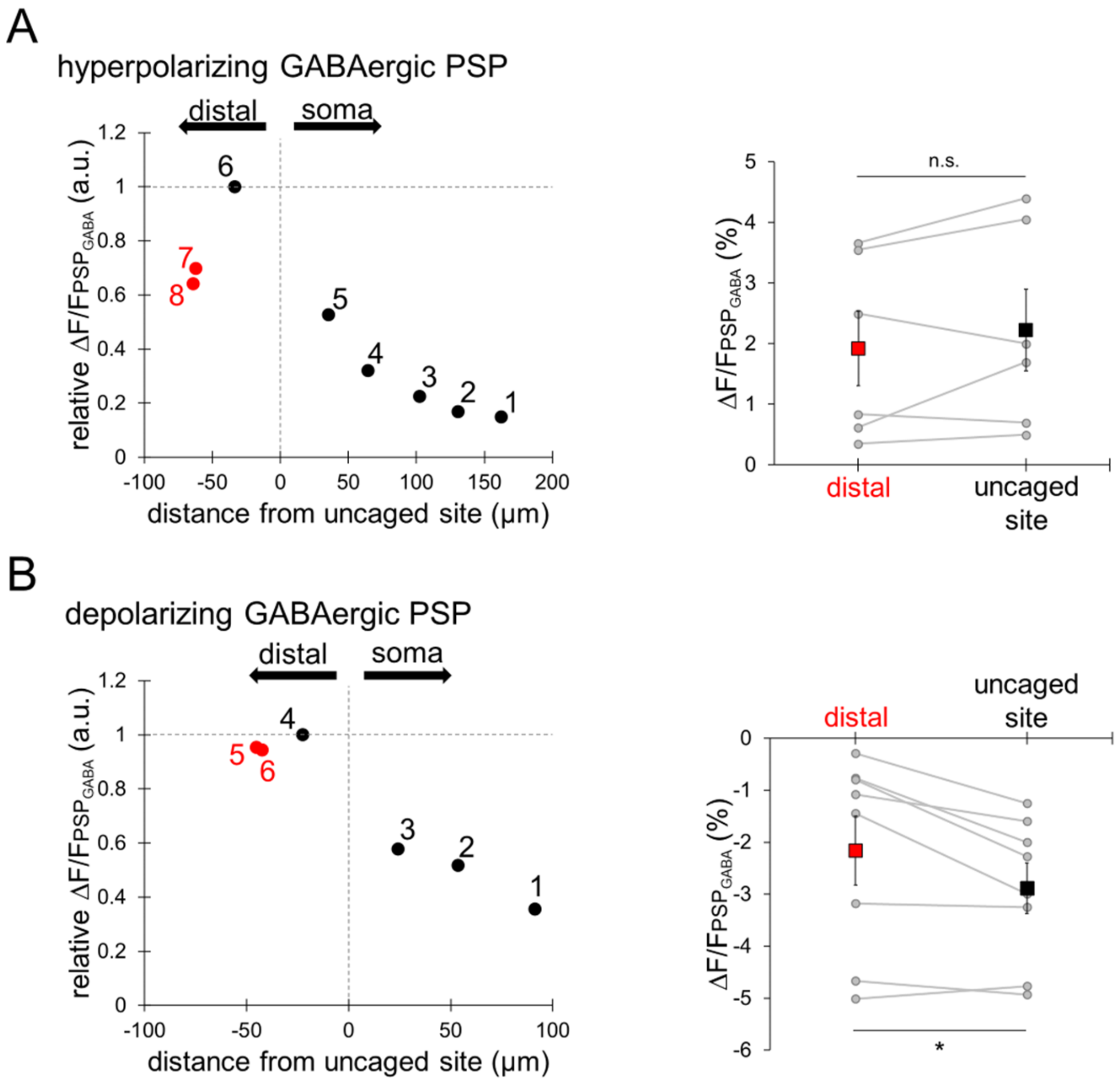


**Fig. S3. Symmetrically attenuating propagation of GABAergic PSPs**

**(A) and (B)** Left, ΔF/F sizes for GABAergic PSPs recorded in a cell shown in Fig. 2A (A) and that in Fig. 2B (B) are plotted against the distance from the uncaged site. Responses at individual ROIs are normalized to the ΔF/F peak at ROI 6 for A or ROI 4 for B (near the uncaged site). Right, absolute ΔF/F for GABAergic PSP at the distal area and that at the uncaged site (A, n = 6 cells, paired t-test, n.s.: not significant; B, n = 8 cells, paired t-test, *: $P < 0.05$). Square plots indicate averages (± SEM).

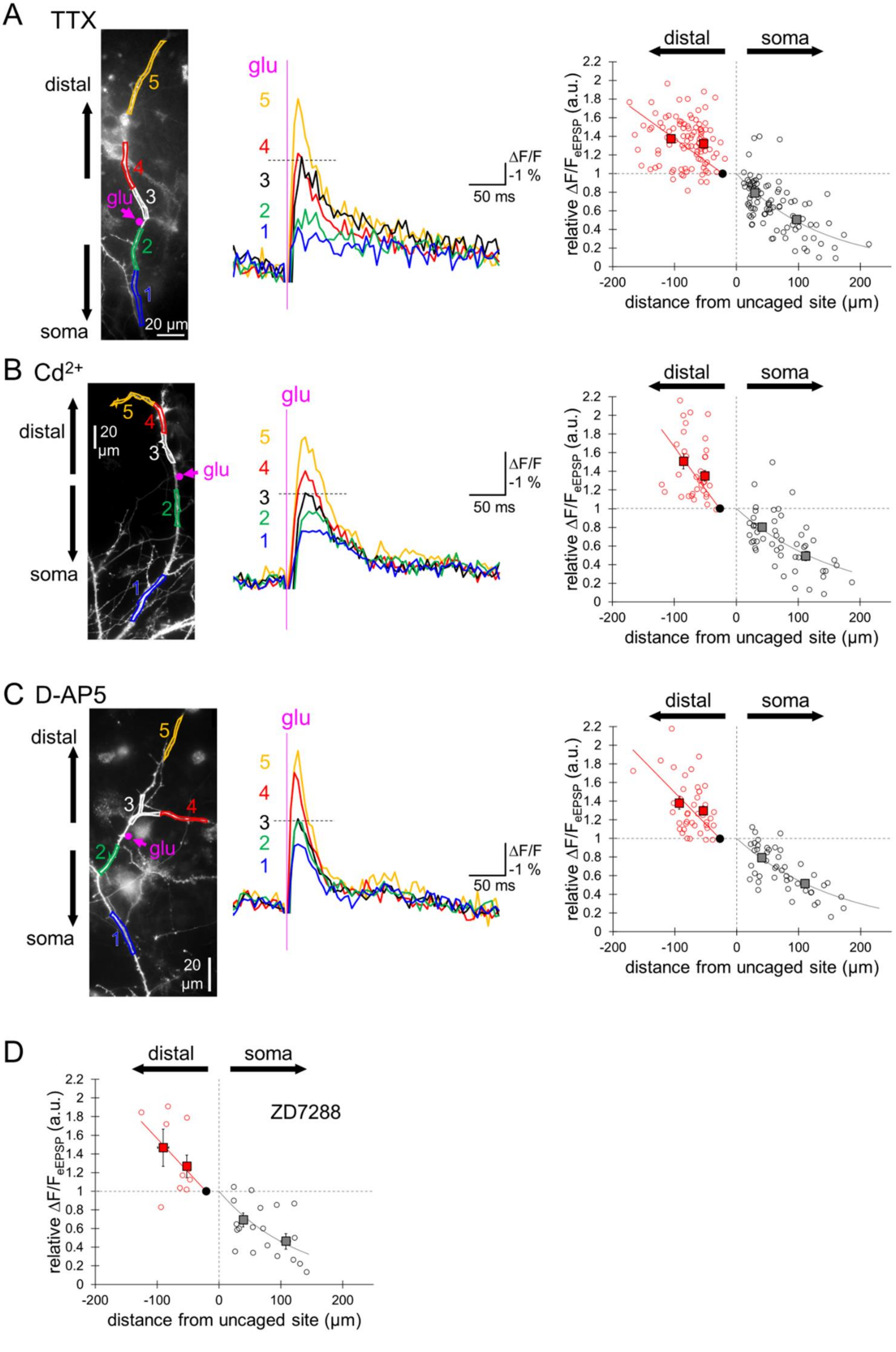

A
TTX
distal
soma
glu
20 µm
5
4
3
2
1
ΔF/F
-1 %
50 ms
distal
soma
relative ΔF/F$_{eEPSP}$ (a.u.)
distance from uncaged site (µm)
B
Cd$^{2+}$
distal
soma
20 µm
glu
5
4
3
2
1
ΔF/F
-1 %
50 ms
distal
soma
relative ΔF/F$_{eEPSP}$ (a.u.)
distance from uncaged site (µm)
C
D-AP5
distal
soma
glu
20 µm
5
4
3
2
1
ΔF/F
-1 %
50 ms
distal
soma
relative ΔF/F$_{eEPSP}$ (a.u.)
distance from uncaged site (µm)
D
distal
soma
ZD7288
relative ΔF/F$_{eEPSP}$ (a.u.)
distance from uncaged site (µm)

**Fig. S4. No abolishment of EPSP amplification by blocking types of cation channel**

**(A-C)** Left, stimulated dendritic segment in the presence of TTX (A, 1 μM), $Cd^{2+}$ (B, 100 μM), or D-AP5 (C, 100 μM). Magenta points indicate the locations of glutamate uncaging. Middle, $\Delta F/F_{eEPSP}$ traces at five ROIs (1-5) indicated in left. Right, relative $\Delta F/F_{eEPSP}$ plotted against the distance from the uncaged site (TTX, 20 cells, red: 103, black: 93 regions; $Cd^{2+}$, 15 cells, red: 37, black: 50 regions; D-AP5, 14 cells, red: 42, black: 45 regions). Square plots indicate averages (± SEM).

**(D)** Relative $\Delta F/F_{eEPSP}$ plotted against the distance from the uncaged site in the presence of 100 μM ZD7288 (6 cells, red: 10, black: 19 regions). Square plots indicate averages (± SEM).

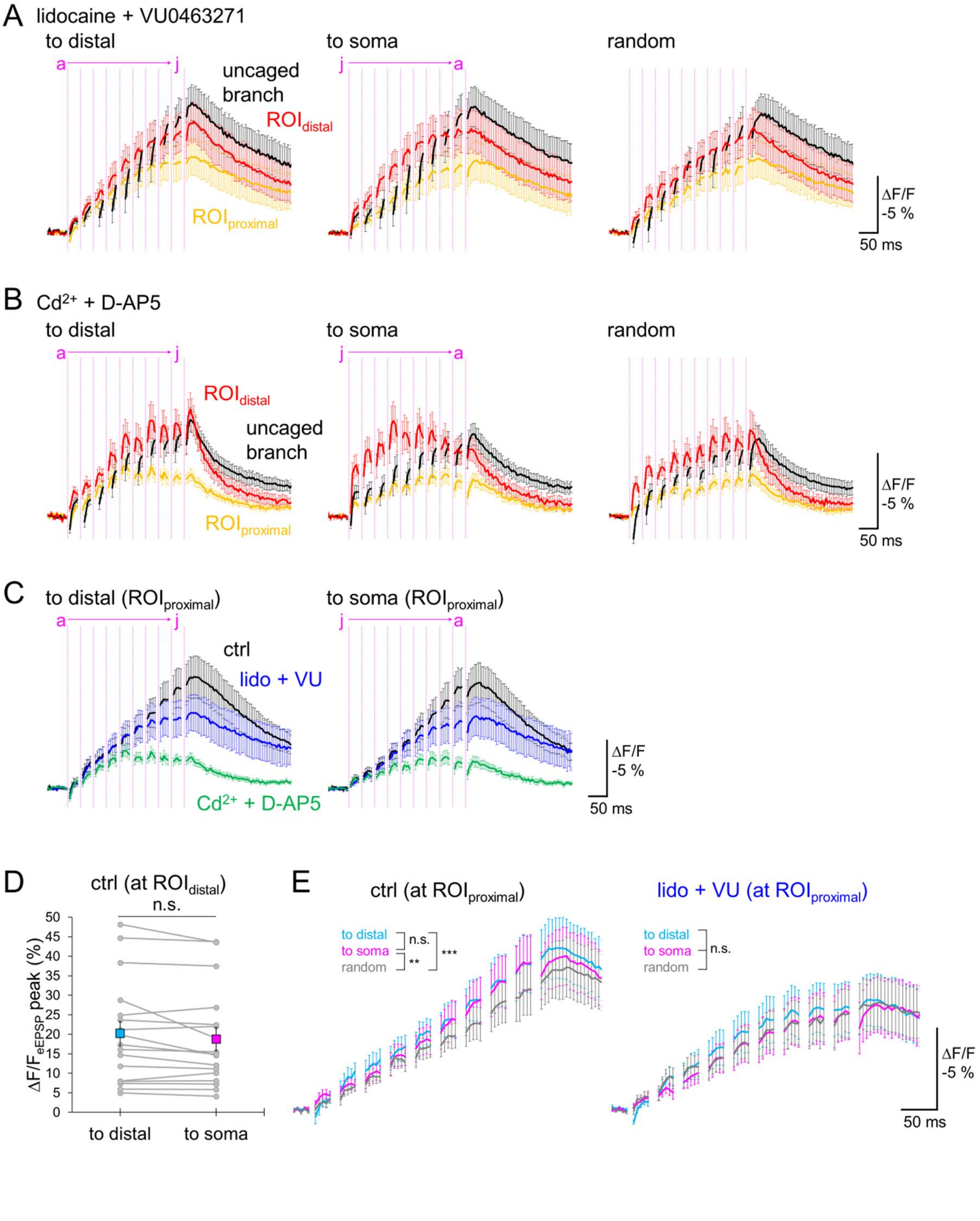

A lidocaine + VU0463271
to distal
to soma
random
a
j
uncaged branch
ROIdistal
ROIproximal
ΔF/F -5 %
50 ms
B Cd2+ + D-AP5
to distal
to soma
random
ROIdistal
uncaged branch
ROIproximal
ΔF/F -5 %
50 ms
C to distal (ROIproximal)
to soma (ROIproximal)
ctrl
lido + VU
Cd2+ + D-AP5
ΔF/F -5 %
50 ms
D ctrl (at ROIdistal)
n.s.
ΔF/FeEPSP peak (%)
0
5
10
15
20
25
30
35
40
45
50
to distal
to soma
E ctrl (at ROIproximal)
to distal
to soma
random
n.s.
**
***
lido + VU (at ROIproximal)
n.s.
ΔF/F -5 %
50 ms

**Fig. S5. Dynamic spatio-temporal integration of synaptic inputs array**

**(A) and (B)** $\Delta F/F_{eEPSP}$ traces (mean ± SEM) at three ROIs (uncaged branch: black, proximal: yellow, distal: red) in three patterns of glutamate uncagings in the presence of 5 mM lidocaine & 10 μM VU0463271 (A, averages of 13 cells), or 100 μM $Cd^{2+}$ & 100 μM D-AP5 (B, 14 cells). Magenta lines indicate 405 nm laser spot illumination.
**(C)** $\Delta F/F_{eEPSP}$ traces (mean ± SEM) at $ROI_{proximal}$ in three conditions: control (black, 17 cells), with lidocaine & VU0463271 (blue, 13 cells), and with $Cd^{2+}$ & D-AP5 (green, 14 cells).
**(D)** $\Delta F/F_{eEPSP}$ peaks at $ROI_{distal}$ in the control condition upon two patterns of laser illumination, "to distal" and "to soma" (n = 17 cells, n.s.: not significant, paired t-test). Square plots indicate averages (± SEM).
**(E)** Comparison of synaptic integration at $ROI_{proximal}$ upon three patterns of glutamate inputs in the absence (ctrl, left) or presence of lidocaine & VU0463271 (right). Two-way ANOVA, **: $P < 0.01$, ***: $P < 0.001$, n.s.: not significant.

| | | #0(soma) | #1~#4 | #5~#7 | #8~#18 | |
|---|---|---|---|---|---|---|
| geometry | L | 30 | 50 | 50 | 30 | [μm] |
| | diam | 30 | 2.5, 2, 1.5, 1 | 0.5 | 0.5 | [μm] |
| electrical property | $C_m$ | 0.8 | 0.8 | 0.8 | 0.8 | [μF/cm$^2$] |
| | $R_a$ | 250 | 250 | 250 | 250 | [Ω·cm] |
| ion | $[Na^+]_{in}$ | 10 | 10 | 10 | 10 | [mM] |
| | $[Na^+]_{out}$ | 145 | 145 | 145 | 145 | [mM] |
| | $[K^+]_{in}$ | 145 | 145 | 145 | 145 | [mM] |
| | $[K^+]_{out}$ | 5 | 5 | 5 | 5 | [mM] |
| | $[Cl^-]_{in}$ | 5 | 5 | 5 | 5 | [mM] |
| | $[Cl^-]_{out}$ | 150 | 150 | 150 | 150 | [mM] |
| leak | $g_{leak.Na}$ | $6.50 \times 10^{-6}$ | $6.50 \times 10^{-6}$ | $6.50 \times 10^{-6}$ | $2.50 \times 10^{-6}$ | [S/cm$^2$] |
| | $g_{leak.K}$ | $3.00 \times 10^{-5}$ | $3.00 \times 10^{-5}$ | $3.00 \times 10^{-5}$ | $2.20 \times 10^{-5}$ | [S/cm$^2$] |
| | $g_{leak.Cl}$ | $2.88 \times 10^{-5}$ | $2.88 \times 10^{-5}$ | $2.88 \times 10^{-5}$ | - | [S/cm$^2$] |
| ClC-2 like channel | $gbar_{Cl}$ | - | - | - | $8.30 \times 10^{-5}$ | [S/cm$^2$] |
| | $V_h$ | - | - | - | -10.2 | [mV] |
| | $V_s$ | - | - | - | 2.5 | [mV] |
| $Na^+/K^+$ pump | $I_{NaKmax}$ | 0.005 | 0.005 | 0.005 | 0.005 | [mA/cm$^2$] |
| | $[ATP]_{in}$ | 5 | 5 | 5 | 5 | [mM] |
| | T | 300.16 | 300.16 | 300.16 | 300.16 | [K] |
| KCC2 | $U_{KCC2}$ | 0.01 | 0.01 | 0.01 | 0.01 | [mA/cm$^2$] |
| Nav | $gbar_{Nav}$ | $3.50 \times 10^{-5}$ | $3.50 \times 10^{-5}$ | $3.50 \times 10^{-5}$ | $3.50 \times 10^{-5}$ | [S/cm$^2$] |

**Table S1. Parameters of the biophysical model**
